# *Legionella pneumophila* LegC7 effector protein drives aberrant ER:endosome fusion in yeast

**DOI:** 10.1101/2020.11.20.391441

**Authors:** Nathan K. Glueck, Kevin M. O’Brien, Vincent J. Starai

## Abstract

*Legionella pneumophila* is a facultative intracellular bacterial pathogen, causing the severe form of pneumonia known as Legionnaires’ disease. *Legionella* actively alters host organelle trafficking through the activities of ‘effector’ proteins secreted via a TypeIVB secretion system, in order to construct the bacteria-laden Legionella-containing vacuole (LCV) and prevent lysosomal degradation. The LCV is derived from membrane derived from host ER, secretory vesicles, and phagosomes, although the precise molecular mechanisms that drive its synthesis remain poorly understood. In an effort to characterize the *in vivo* activity of the LegC7/YlfA SNARE-like effector protein from *Legionella* in the context of eukaryotic membrane trafficking in yeast, we find that LegC7 interacts with the Emp46p/Emp47p ER-to-Golgi glycoprotein cargo adapter complex, alters ER morphology, and induces aberrant ER:endosome fusion, as measured by visualization of ER cargo degradation, reconstitution of split-GFP proteins, and enhanced oxidation of the ER lumen. LegC7-dependent toxicity, disruption of ER morphology, and ER:endosome fusion events were dependent upon endosomal VPS class C tethering complexes and the endosomal t-SNARE, Pep12p. This work establishes a model in which LegC7 functions to recruit host ER material to the bacterial phagosome during infection by inducing membrane fusion, potentially through interaction with host membrane tethering complexes and/or cargo adapters.

## Introduction

*Legionella pneumophila* is a ubiquitous Gram-negative, facultatively intracellular bacterium, capable of invading and replicating within a broad range of freshwater protozoa[1, 2]. Upon aerosolization of contaminated water supplies and subsequent inhalation by susceptible individuals, *Lpn* is taken up into alveolar macrophages via coiling phagocytosis[3], where the bacterium persists and replicates within a modified membrane-bound compartment termed the *Legionella*-containing vacuole (LCV). Left untreated, this infection can lead to the severe form of pneumonia known as Legionnaires’ disease, which can have a mortality rate of at least 10%[4]. Localized outbreaks of legionellosis continue to rise worldwide, and the number of reported cases in the United States in 2017 was ~7,500, an increase of more than 500% when compared to cases reported in 2000[5]. Like most intracellular bacteria, *L. pneumophila* actively manipulates host physiology in order to create its replicative niche. The *L. pneumophila* genome encodes for a Dot/Icm type-IVB secretion system, which secretes over 300 confirmed and predicted “effector” proteins into the host cell upon contact and internalization[6]. These effectors are known to modulate myriad host cell processes to ensure the survival of the bacterium, including: protein and membrane trafficking pathways[7, 8], autophagy[9], and actin dynamics[10]; these activities are coordinated to prevent the fusion of the *Legionella*-containing phagosome with endolysosomal degradative compartments and recruit ER-derived vesicles to the phagosome to generate the LCV[11]. Understanding the mechanisms by which these effectors can modulate eukaryotic physiology has been a focus of intense study[6, 12], but which has been complicated by the fact that activities of these proteins are thought to be highly redundant, as few single effector gene disruptions lead to pathogenicity defects in model systems[12]. Therefore, the use of heterologous eukaryotic expression systems to dissect the activities of individual *Lpn* effectors has proven fruitful[7, 8, 13, 14].

LegC7/YlfA is a Dot/Icm protein substrate that was originally identified as a *L. pneumophila* effector that inhibits growth of *Saccharomyces cerevisiae* when expressed, although the mechanism of toxicity was not determined[14]. When expressed in CHO cells, it was found that LegC7/YlfA localized to ER and early secretory vesicles, and that this localization was dependent upon its N-terminal hydrophobic/transmembrane domain[14]. Furthermore, LegC7 expression in yeast resulted in general vacuolar protein sorting defects[7], and specifically inhibited endosomal cargo delivery to the degradative vacuole; other protein trafficking pathways to the vacuole were not affected[15]. Interestingly, LegC7/YlfA and two other *Lpn* effectors, LegC2/YlfB (a paralog of LegC7) and LegC3, have been described as eukaryotic “SNARE-like” proteins, as they have similar coiled-coil domain structures and can be found to engage in higher-order complexes, similar to the SNARE proteins responsible for catalyzing the majority of eukaryotic intracellular membrane fusion events[16]. Indeed, Δ*legC7* Δ*legC2* mutant strains of *Legionella* form aberrant LCVs during infection that contain less host ER-derived membrane, suggesting a role for LegC7 and/or LegC2 in ER membrane recruitment or fusion to the phagocytic membrane surrounding *Legionella*. Despite the reported importance of LegC7 and LegC2 in the synthesis of the LCV, these Δ*legC7*Δ*legC2* mutant strains do not display proliferation defects during pathogenesis[17]. Biochemical evidence of the SNARE-like membrane fusion activity of complexes containing LegC2, LegC7, and LegC3 was discovered when synthetic liposomes harboring LegC2/LegC7/LegC3 “Q-SNARE” complexes were fused with target liposomes harboring a host endosomal R-SNARE, VAMP4[18]; this SNARE complex activity was also found to be completely resistant to the activity of the conserved SNARE complex remodeling complex, NSF/α-SNAP[18, 19]. Therefore, it is extremely likely that the LegC proteins are utilized by *L. pneumophila* to recruit host membrane compartments, including ER and VAMP4-containing vesicles to the LCV during pathogenesis via hijacking host SNARE machineries.

Based on these previous works, we set out to reconcile the observation that *L. pneumophila* LegC7 is toxic to yeast with its ability to both interact with ER and endosomal compartments in model host cells and alter endosomal membrane trafficking pathways in yeast. We now find that high-level expression of LegC7 induces strong ER morphology defects in yeast, creating aberrant ER membrane structures that also colocalize with Vps8p-containing endosomal compartments. Both the ER morphology defects and growth inhibition observed upon LegC7 expression are completely reversed by deletions in so-called “class C” complex core genes (*VPS11, VPS16, VPS18, VPS33*) which comprise the core subunits of two endolysosomal multisubunit tethering complexes, CORVET (class C core vacuole/endosome tethering) and HOPS (homotypic fusion and vacuole protein sorting). We also observe that LegC7 expression increases the oxidation state of the ER lumen and induces the reconstitution of split-GFP protein fragments contained within ER and endosomal compartments. Coupled with the finding that LegC7 induces the degradation of the ER lumenal Kar2p protein in a vacuolar protease-dependent manner, we now provide evidence that LegC7 directs the fusion of ER-derived compartments with endosomes in a CORVET/HOPS-dependent manner, thereby bypassing normal ER:Golgi trafficking. These results provide additional evidence that LegC proteins from *Legionella* directly recruit ER membrane to the LCV during infection.

## Results

### LegC7 immunoprecipitates an ER:Golgi cargo adapter complex

LegC7/YlfA expression in yeast is known to be toxic[14], although the expression of the similar coiled-coil secreted effectors from *Legionella* (LegC2/YlfB and LegC3) is not toxic under the same growth conditions[13, 14]; LegC7 therefore appears to have an in vivo activity distinct to that of LegC2/3. In an effort to elucidate the mechanism of LegC7-mediated growth inhibition, whole cell extracts were generated from cells expressing LegC7 or empty vector controls, incubated with Protein A:α-LegC7 resin (Materials and Methods), and resultant immunoprecipitates were subjected to LC-MS/MS for total protein identification. When eliminating proteins identified across both pulldown conditions, we noted a strong enrichment of the ER:Golgi glycoprotein cargo adapters, Emp47p and Emp46p (**Table 1**)[20, 21]. In addition, we noted the enrichment of the Emp47p-interacting protein, Ssp120p, suggesting a possible interaction of LegC7 with, or enrichment within, ER-derived COPII-coated vesicles.

**Table 1.**
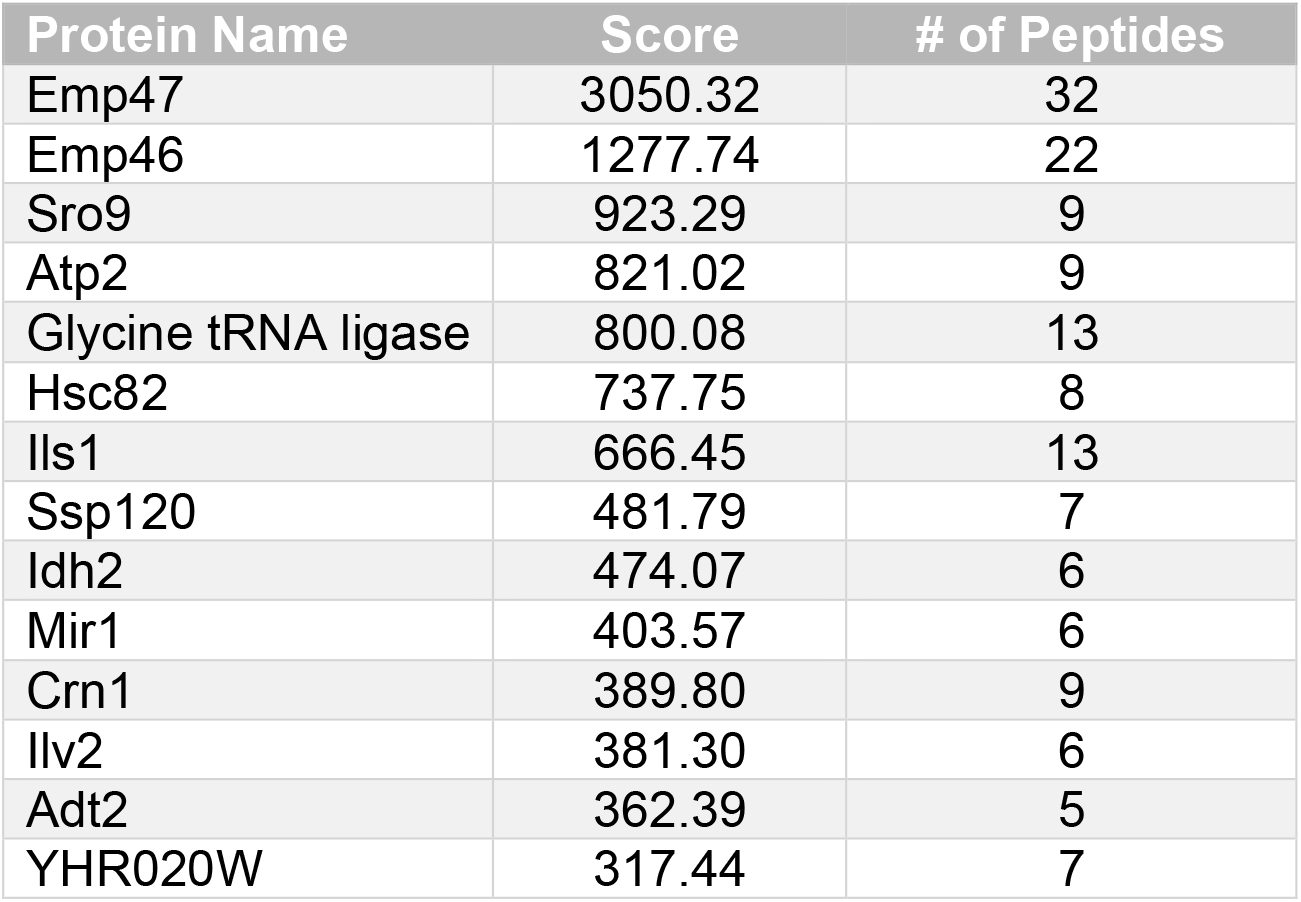
Protein ID list from LegC7 immunoprecipitations (scores > 300).

As LegC7 appears to interact with an Emp47p-containing complex, we considered the possibility that LegC7 was glycosylated in vivo. To address this, we purified LegC7 from yeast extracts using the same Protein A:α-LegC7 resin used previously. SDS-PAGE bands containing LegC7 were excised and submitted to the University of Georgia Complex Carbohydrates Research Center for *N*-glycan analysis via nano-LC-MS/MS (Supplemental Materials and Methods). While peptides from LegC7 were clearly detected (**Fig. S1, Table S3**), no glycosylations or other post-translational modifications on any LegC7 peptide were discovered (**Fig. S2A-R**). Therefore, while LegC7 appears to interact with a known glycoprotein cargo receptor, this interaction either does not require glycosylation of LegC7, or this interaction is an indirect one.

### *EMP46/47* deletions mimic the LegC7-induced inhibition of carboxypeptidase S trafficking

Given the apparent interaction between the Emp47p complex and LegC7, as well as our previous observation that LegC7 expression inhibits the delivery of GFP-tagged carboxypeptidase S (CPS-GFP) to vacuoles[15], we considered the possibility that LegC7 may interfere with Emp47p complex function, which then results in the previously-observed CPS-GFP trafficking defects. Therefore, we observed the CPS-GFP trafficking phenotypes of *emp46*Δ *emp47*Δ strains and compared them to wild type strains expressing LegC7. Wild type yeast cells expressing CPS-GFP show a vacuolar lumenal distribution with few cytosolic punctae in each cell, likely representing endosomes containing CPS-GFP (**Figs. 1A,C**, **and** **D**). Expression of LegC7 results in the accumulation of punctate structures and far fewer cells containing vacuolar CPS-GFP, as seen previously[15] (**Figs. 1C** **and** **D**). When CPS-GFP localization is observed in an *emp46*Δ *emp47*Δ strain, we noted that fewer cells showed vacuolar localization of CPS-GFP, with a corresponding increase in cytosolic punctae, similar to that observed in LegC7-expressing strains (**Figs. 1B-D**). As the cellular distribution of CPS-GFP is overall similar between LegC7-expressing and Emp-deficient strains, it is likely that LegC7 expression interferes with Emp47p complex function in some manner. Interestingly, however, *emp46*Δ *emp47*Δ deletion strains do not strongly reduce the LegC7-mediated growth inhibition (**Fig. S3**), suggesting that LegC7 does not absolutely depend upon the presence of Emp46/47 for function in vivo. Given the fact that Emp47p and Emp46p form an important cargo adapter complex residing in the ER and on COPII vesicles, however, we next decided to observe the effects of LegC7 expression on ER morphology.

**Figure 1.**
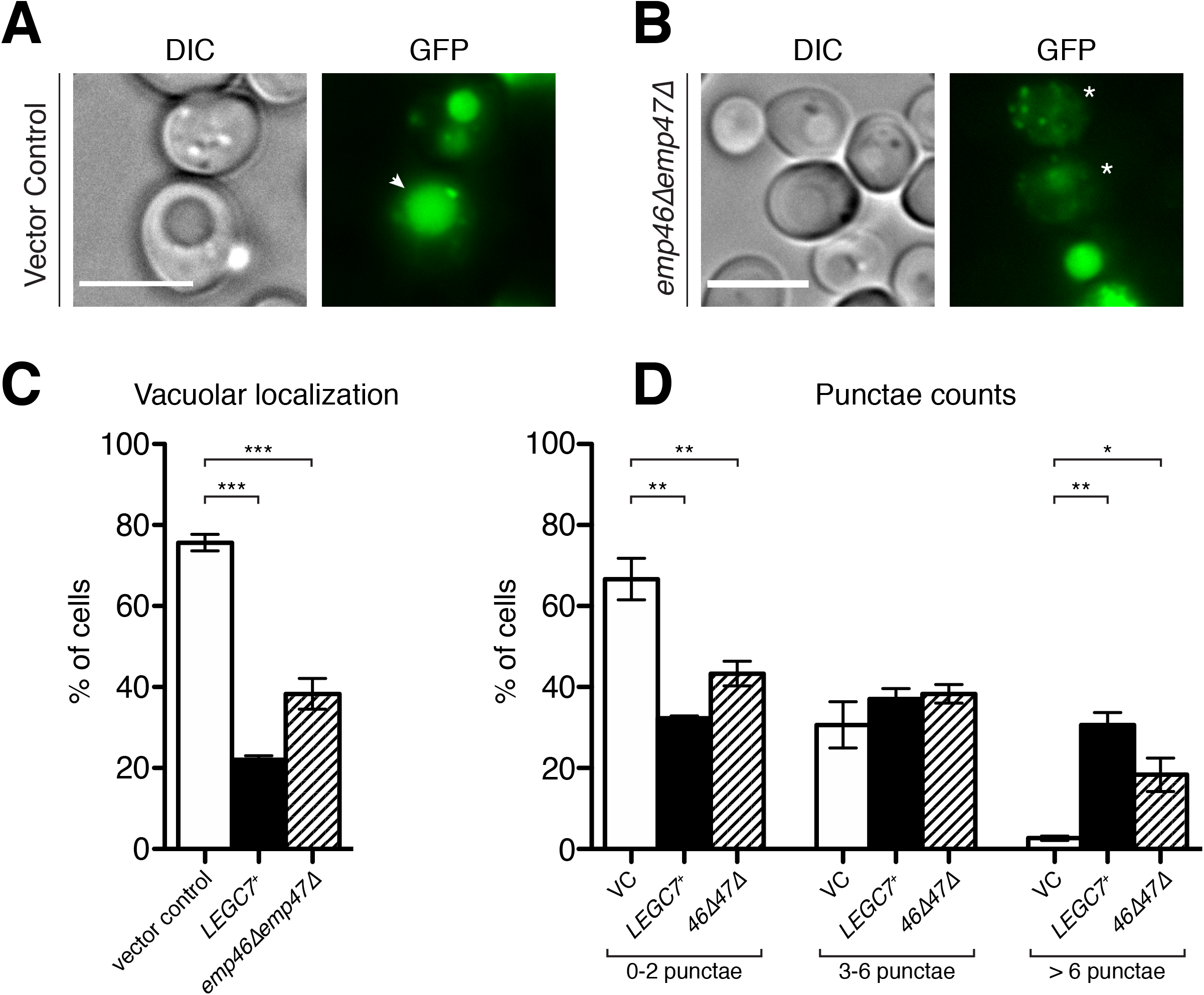
An *emp46*Δ*emp47*Δ double mutation and LegC7 expression both alter localization of Carboxypeptidase S. Yeast strains harboring GFP-CPS were visualized to determine vacuolar localization and number of punctae. All strains were grown to saturation in CSM medium at 30°C, harvested via centrifugation, resuspended in an equal volume of fresh CSM containing 1% raffinose and 1% galactose, and grown for an additional 18 h at 30°C before imaging. (**A**) and (**B**) are representative images of cells with and without vacuolar localization, respectively. The white arrow indicates vacuolar localization and the white astrisks indication cells without vacuolar localization. Scale bars = 5 μ. Individual cells were analyzed to determine vacuolar localization of GFP (**C**) and number of punctae (**D**). ≥ 100 cells each per three independent experiments were analyzed; error bars represent ± the standard deviation between experiments; (*) = P < 0.05; (**) = P < 0.005; (***) = P < 0.0005.

### LegC7 expression alters normal ER morphology

Based upon the discovery that LegC7 immunoprecipitates proteins known to reside in COPII-coated vesicles (Emp46p, Emp47p, and Ssp120p) involved in ER:Golgi protein transport, we decided to observe the effects of LegC7 expression on gross ER and Golgi morphologies. Yeast strains expressing GFP fusions of an ER translocon subunit, Sec63p, show the expected distribution of perinuclear and cortical ER membrane (**Fig. 2A**)[22]. Expression of a fluorescently-tagged LegC7, LegC7-mRuby2, however, induces a drastic redistribution of perinuclear ER membrane in cells which clearly express LegC7-mRuby2 (**Fig. 2A**). Strikingly, LegC7-mRuby2 appeared to accumulate in large punctate structures that overlapped with large accumulations of Sec63-positive membranes (**Fig. 2A**, **arrowheads**); these large Sec63-GFP accumulations were not observed in vector control strains. To ensure that the LegC7-mRuby2 fusion protein was still active in vivo, we confirmed that strains expressing LegC7-mRuby2 were inhibited for growth to the same extent as untagged LegC7 (**Fig. S4A**); N-terminal, GFP-tagged LegC7 was no longer toxic (**Fig. S4A**), even when strains expressed nearly equivalent levels of LegC7 (**Fig. S4B**). Therefore, we concluded that LegC7-mRuby2 still maintained LegC7 activity, and this activity results in ER morphology defects upon expression. To determine whether LegC7 expression also disrupts Golgi morphology, we performed similar experiments in strains expressing GFP-Vrg4, a *cis*-Golgi GDP-mannose transporter[23]. In control cells, *cis*-Golgi structures were observed to be generally punctate, with some perinuclear localizations (**Fig. 2B**), in agreement with previously reported structures[23]. In the presence of LegC7-mRuby2 expression, we noted similar punctate and perinuclear structures, and LegC7-mRuby2 did not strongly colocalize with any Vrg4-GFP-positive membranes (**Fig. 2B**). Taken together, these images suggest that LegC7 expression does not strongly alter normal cis-Golgi morphology, while having strong effects on normal ER morphology.

**Figure 2.**
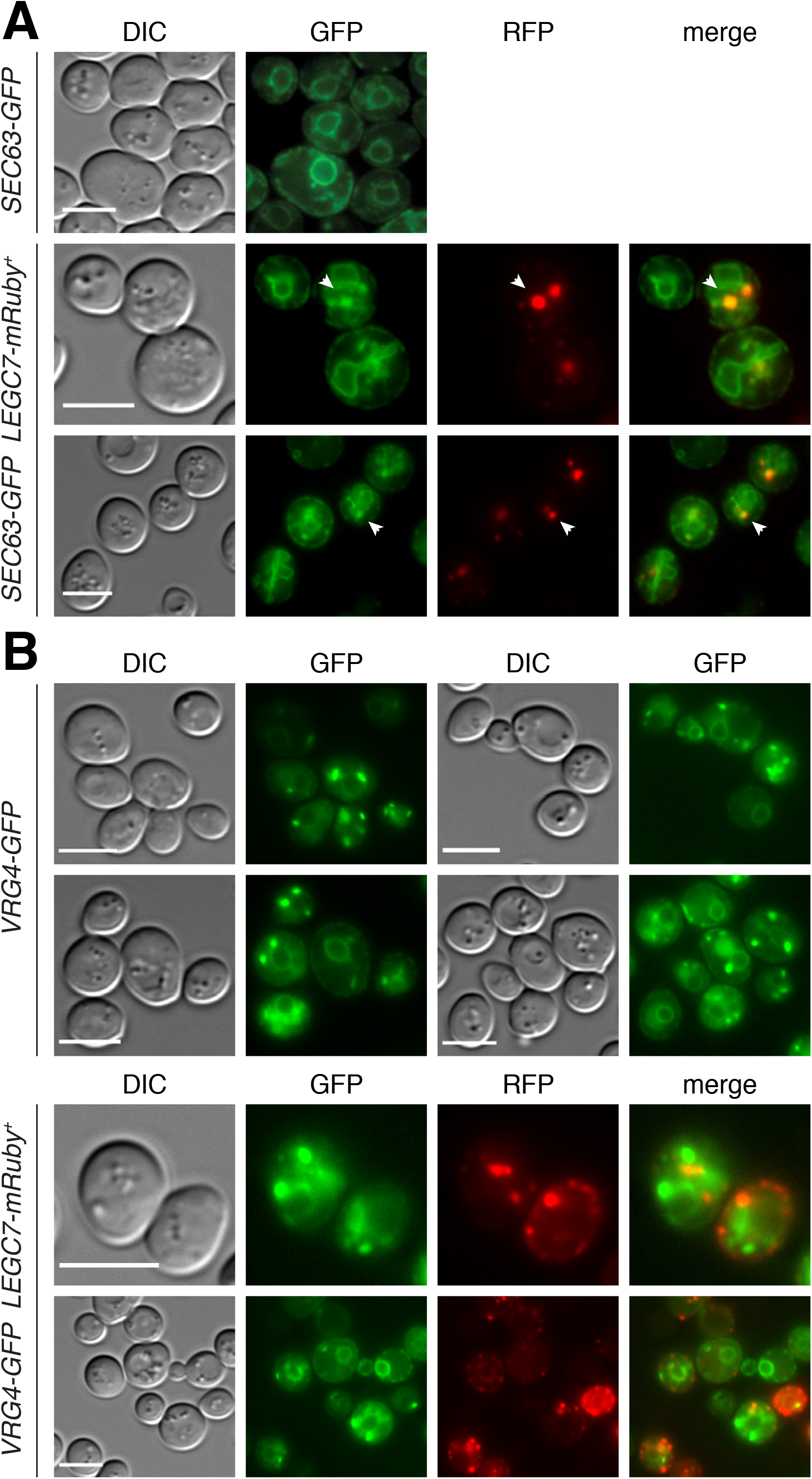
LegC7 alters ER morphology but doesn’t affect Golgi morphology in a discernable manner. Yeast strains expressing LegC7-mRuby2 and GFP fusions of ER translocon subunit Sec63p (**A**) or cis-Golgi GDP-mannose transporter Vrg4p (**B**) were visualized. All strains were grown to saturation in CSM medium at 30°C, harvested via centrifugation, resuspended in an equal volume of fresh CSM containing 1% raffinose and 1% galactose, and grown for an additional 18 h at 30°C before imaging. Arrows indicate patches where LegC7-mRuby2 and Sec63-GFP colocalize. Scale bars = 5 μ.

### Deletion of the VPS class C tethering complex subunits suppresses LegC7-mediated growth inhibition

In parallel to observing the effects of LegC7 expression on ER membrane structure, we also considered that LegC7 expression was previously shown to inhibit the delivery of proteins to the degradative vacuole via both biosynthetic and endocytic pathways[15]. To identify the potential targets of LegC7 activity on potential endolysosomal or protein trafficking targets, we performed a directed screen of 86 gene deletions that focused on general membrane trafficking mechanisms to identify strains that continued to grow in the presence of LegC7 expression (**Table S2**). Among all gene deletions screened, deletions of the CORVET and HOPS tethering core complex subunits (Vps11p, Vps16p, Vps18p, Vps33p) resulted in nearly complete restoration of growth in the presence of LegC7 expression (**Fig. 3A**). CORVET (class C core vacuole/endosome tethering) and HOPS (homotypic fusion and protein sorting) are hexameric tethering complexes that facilitate the SNARE-dependent fusion of endolysosomal compartments through direct interactions with Rab-family GTPases, SNARE complexes, and membrane lipids across apposed organelles[24, 25].

**Figure 3.**
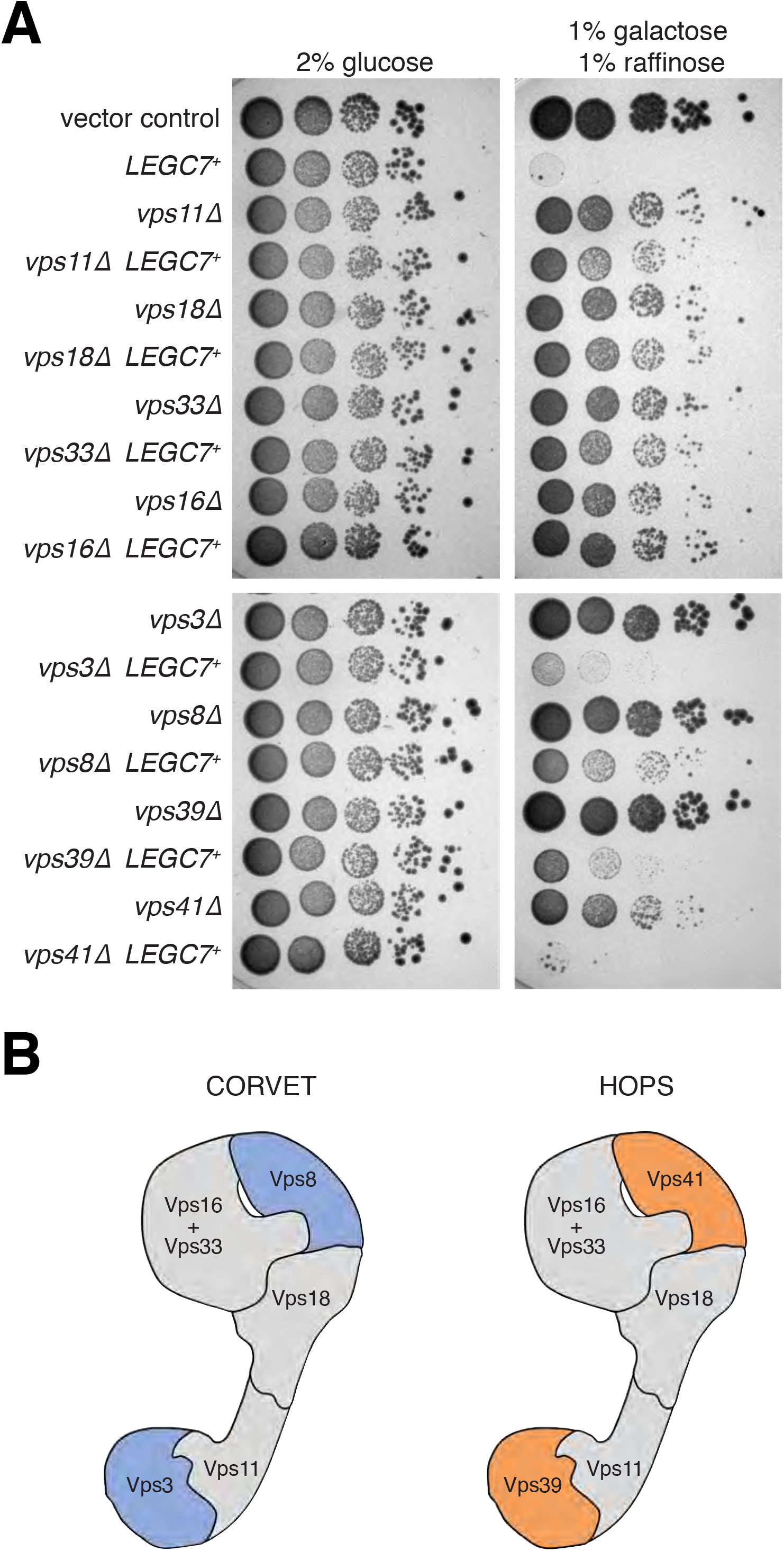
Deletion of Class C tethering complex subunits suppresses LegC7-mediated growth inhibition. (**A**) Yeast deletion mutants harboring either pYES2NT C or pYES2-*LEGC7*^+^ were grown to saturation in CSM at 30°C. For each strain, 1 OD600 unit was harvested via centrifugation, resuspended in 1 mL of 0.9% NaCl, and serially diluted 1:10 four times. 5 μL of each dilution was plated onto CSM containing 2% glucose and CSM containing 1% galactose and 1% raffinose to induce LegC7 expression. (**B**) Models of the class C tethering complexes are displayed, with the core complex in gray and the Rab-specific subunits in blue and orange.

In addition to the shared core subunits listed above, each distinct tethering complex contains two unique and interchangeable subunits, which mediate binding specificity to Rab-family GTPases (**Fig. 3B**). CORVET contains Vps3p and Vps8p, which bind to the Rab5 homolog, Vps21p, found on early endosomal membranes[26, 27]. HOPS complex contains Vps39p and Vps41p, which bind to the Rab7 homolog, Ypt7p, found on late endosomal and vacuolar membrane compartments[28-30]. As both HOPS and CORVET complexes are disrupted in class C core mutant strains, we individually deleted each of the Rab-specific subunits of these complexes to determine the effects of CORVET- or HOPS-specific disruptions on LegC7 activity. Unlike what we observed with the core subunit deletions, however, deletions of the Rab-specific subunits resulted in varying degrees of restored growth in the presence of LegC7 expression **(Fig. 3A)**. It was interesting to note that, of the four interchangeable subunits, deletions of the CORVET-specific *VPS8* gene provided the strongest restoration of growth to LegC7-expressing strains under these conditions. As Vps8p interacts directly with the endosomal Rab GTPase, Vps21p, as a part of the CORVET complex[26, 27], it raised the possibility that proper endosomal fusion dynamics is required for the *in vivo* toxicity of LegC7 in yeast.

### LegC7 induces colocalization of Vps8p and ER

As class C core gene deletions and *VPS8* deletions appeared to suppress the toxicity of LegC7 expression, we decided to visualize the localization of some representative CORVET/HOPS complex subunits in the presence of LegC7. Vps8-GFP (CORVET) normally forms a punctate pattern in the cell (**Fig. 4A**), as expected[31]. Upon co-expression of LegC7-mRuby2, however, we find Vps8-GFP punctae that strongly colocalize with LegC7-mRuby2 (**Figs. 4A** **and** **4F**). Expression of LegC7 in strains harboring Vps33-GFP, representative of the class C core complex, does not dramatically alter the localization of Vps33-GFP, nor does LegC7 appreciably colocalize with Vps33-GFP/class C core (**Figs. 4B** **and** **4F**). Also, in contrast to the colocalization observed between LegC7 and Vps8p, LegC7-mRuby2 did not colocalize with the endosomal RabGTPase, Vps21, which is a binding partner of the CORVET complex (**Fig. 4D**).

**Figure 4.**
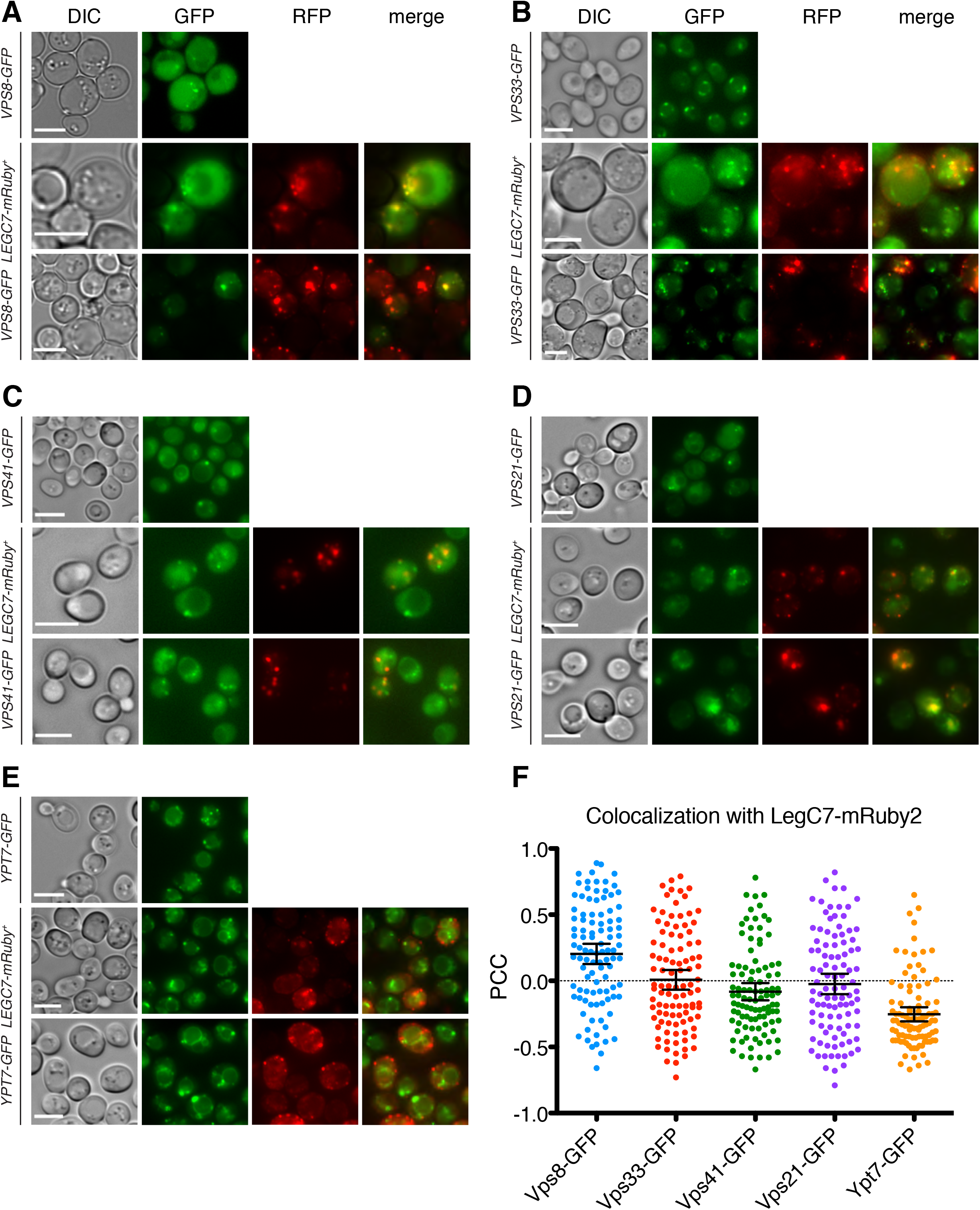
LegC7 colocalizes with early endosomal fusion machinery more than that of late endosomes. Yeast strains expressing LegC7-mRuby2 and GFP fusions CORVET-specific subunit Vps8p (**A**), Class C core complex subunit Vps33p (**B**), HOPS-specific subunit Vps41p (**C**), early endosomal Rab5 homolog Vps21p (**D**), and late endosomal Rab7 homolog Ypt7p (**E**) were visualized. All strains were grown to saturation in CSM medium at 30°C, harvested via centrifugation, resuspended in an equal volume of fresh CSM containing 1% raffinose and 1% galactose, and grown for an additional 18 h at 30°C before imaging. Scale bars = 5 μ. (**F**) Colocalization of GFP and mRuby2 was quantified in the form of Pearson correlation coefficients (PCCs) for 100 cells per strain using the Coloc2 plugin in Fiji (ImageJ). Error bars represent 95% confidence intervals of the average PCC.

Because LegC7-mRuby2 colocalized with both Vps8-GFP and Sec63-GFP punctae (**Fig. 2A**), we then considered the possibility that these Vps8p-positive and Sec63-positive structures also colocalized in response to LegC7. In strains lacking LegC7, Vps8-GFP and Sec63-RFP form expected morphologies and do not colocalize (**Fig. 5A**). In the presence of LegC7, however, we noted both defective ER morphology coupled with a strong recruitment of Vps8-GFP to the Sec63-RFP punctae (**Fig. 5A**); these structures may therefore represent accumulation of endosomal material on the ER membrane. Due to the requirements of CORVET/HOPS subunits for LegC7 toxicity, coupled with the observation that endosomal material accumulates on the ER in a LegC7-dependent manner, we sought to determine if CORVET/HOPS complex activity was required for the LegC7-dependent disruption of ER morphology.

**Figure 5.**
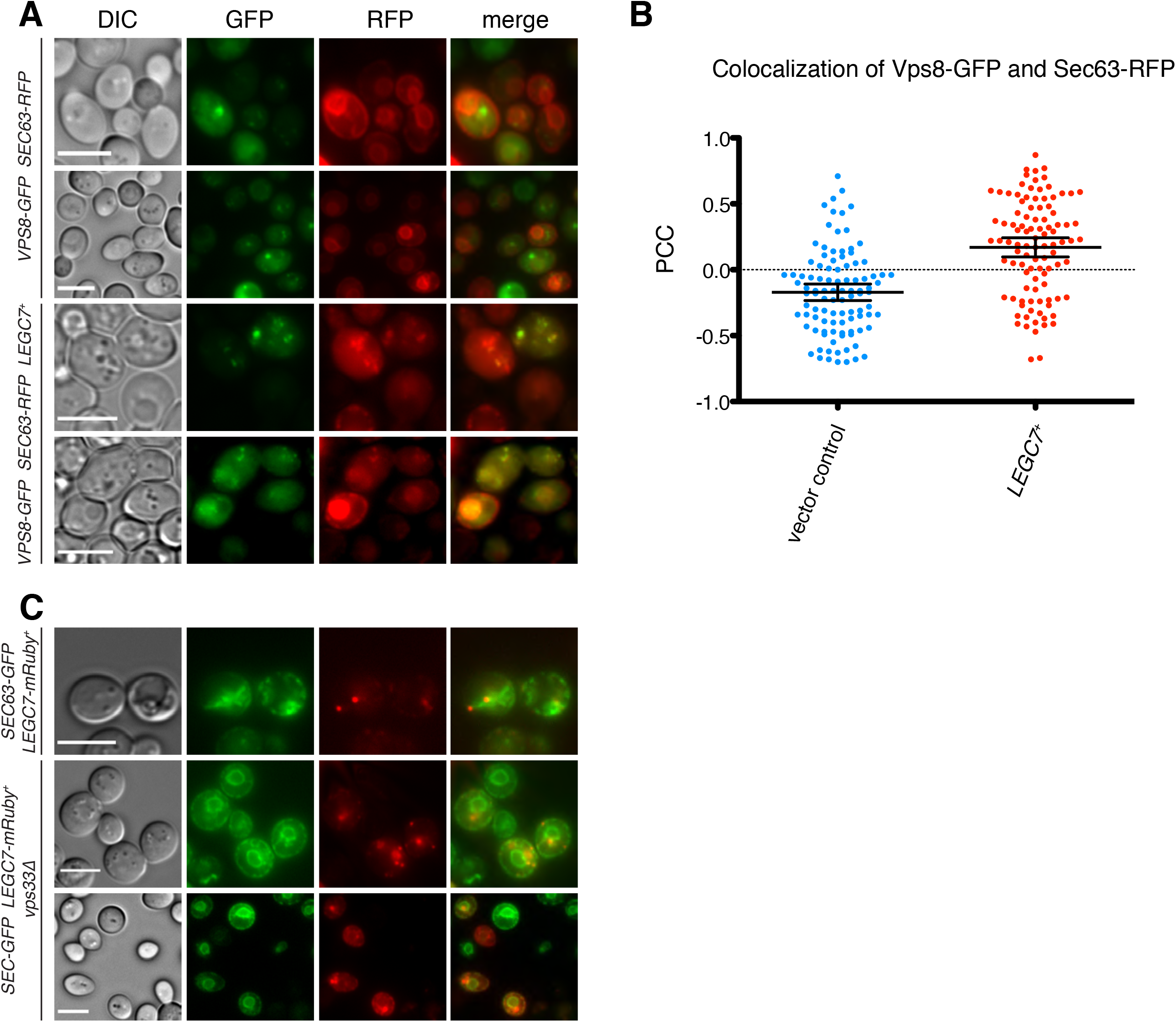
CORVET subunit Vps8p localizes to the ER during LegC7 expression, and endosomal tethering machinery is required for LegC7-induced alteration of ER morphology. (**A**)Yeast strains containing Vps8-GFP, Sec63-RFP, and pYES2NT C or pYES2-*LEGC7*^+^ were visualized. (**B**) Colocalization of GFP and RFP was quantified in the form of Pearson correlation coefficients (PCCs) for 100 cells per strain in (**A**) using the Coloc2 plugin in Fiji (ImageJ). Error bars represent 95% confidence intervals of the average PCC. (**C**) Yeast strains containing Sec63-GFP and expressing LegC7-mRuby2 were visualized to observe the effect of a *vps33*Δ deletion on ER morphology during LegC7 expression. All strains were grown to saturation in CSM medium at 30°C, harvested via centrifugation, resuspended in an equal volume of fresh CSM containing 1% raffinose and 1% galactose, and grown for an additional 18 h at 30°C before imaging. Scale bars = 5 μ.

To address this possibility, both LegC7-mRuby2 and Sec63-GFP were expressed in a *vps33*Δ mutant strain, which lacks functional CORVET/HOPS complexes. Strikingly, the aberrant ER phenotype normally observed in strains expressing LegC7 (**Fig. 5C**) was not present, and the ER morphology of these cells appeared to be completely wild type. Therefore, these data suggest that the function of endosomal tethering complexes plays a role in the aberration of the ER observed with LegC7 expression, despite the fact that CORVET/HOPS complexes function exclusively in endolysosomal membrane dynamics and play no known role in ER-to-Golgi transport[24, 25].

### LegC7 causes the degradation of ER lumenal content

As LegC7 has an N-terminal transmembrane domain[14] and induced abnormal ER and endolysosomal structures upon expression, we attempted to identify the membrane compartment to which LegC7 localized via subcellular fractionation. Total membranes from yeast strains expressing LegC7 were fractionated in a density-dependent manner. Each fraction from the gradient was TCA precipitated and probed via western blotting for ER and Golgi marker proteins and LegC7. While LegC7 appeared to localize to membranes with a density between ER (Kar2p) and Golgi (Och1p) (**Fig. 6A**), we also noted a faint degradation product of the ER lumenal protein Kar2p in the low-density fractions of the LegC7-expressing strain; this product was never observed in any fractions of the control strain (**Fig. 6A**). With this finding, we considered the possibility that this degradation results from the fusion of endolysosomal compartments and ER facilitated by LegC7, thereby exposing Kar2p to vacuolar proteases.

**Figure 6.**
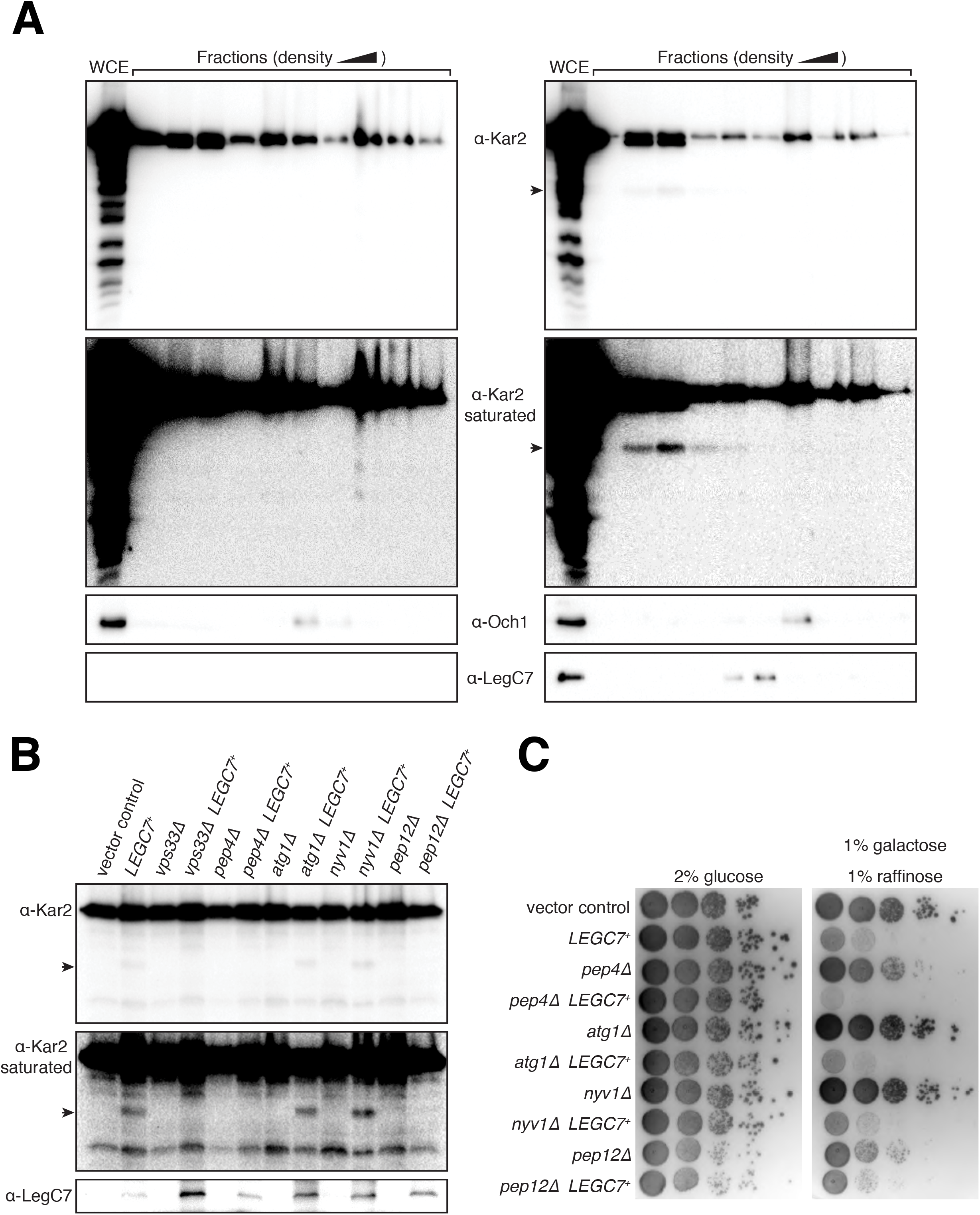
LegC7 causes the degradation of ER luminal ATPase Kar2p, dependent upon endosomal fusion machinery and vacuolar proteases. (**A**) Yeast strains containing pYES2NT C or pYES2-*LEGC7*^+^ were grown to saturation in CSM medium at 30°C, harvested via centrifugation, resuspended in an equal volume of fresh CSM containing 1% raffinose and 1% galactose, and grown for an additional 18 h at 30°C. Equal amounts of each strain were harvested via centrifugation, dounced, fractionated into 10 fractions, and fractions were TCA precipitated (Materials and Methods). Equal volumes of each fraction were separated via SDS-PAGE and immunoblotted for the ER luminal ATPase Kar2p, the cis-Golgi mannosyltransferase Och1p, and LegC7. (**B**) Strains were treated as in (**A**) but not fractionated. Equal volumes of whole cell extracts were separated via SDS-PAGE and immunoblotted for Kar2 and LegC7. The black arrows in (**A**) and (**B**) indicate Kar2 degradation product caused by LegC7. (**C**) Yeast deletion mutants harboring either pYES2NT C or pYES2-*LEGC7*^+^ were grown to saturation in CSM at 30°C. For each strain, 1 OD600 unit was harvested via centrifugation, resuspended in 0.9% NaCl, and serially diluted 1:10 four times. 5 μL of each dilution was plated onto CSM containing 2% glucose and CSM containing 1% galactose and 1% raffinose to induce LegC7 expression.

To test whether LegC7-dependent degradation of Kar2p requires vacuolar proteases, whole cell extracts of a vacuolar protease-deficient *pep4*Δ mutant expressing LegC7 were probed for the Kar2p degradation product. The degradation product does not appear in a *pep4*Δ mutant, demonstrating a requirement for vacuolar proteases for LegC7-dependent Kar2p degradation (**Fig. 6B**). This Kar2p degradation is also absent in both a CORVET/HOPS-deficient *vps33*Δ mutant and an endosomal t-SNARE-deficient *pep12*Δ mutant, demonstrating that endosomal fusion machinery is necessary for this degradation event as well. Notably, LegC7-dependent Kar2p degradation does appear in an *atg1*Δ mutant, confirming that this degradation event is independent of cellular autophagic processes, including ER-phagy[32, 33].

Given that LegC7 causes vacuolar protease-dependent degradation of Kar2p, mutants screened for Kar2p degradation were also screened for LegC7-induced growth inhibition to determine if growth inhibition is a direct result of degradation of ER lumenal contents. Aside from the complete restoration of growth in the *vps33*Δ strain (**Fig. 3A**), *pep12*Δ provides suppression of LegC7 toxicity, while *pep4*Δ does not (**Fig. 6C**), strongly supporting the hypothesis that while endosomal fusion machinery is required for vacuolar protease-dependent degradation of Kar2p, simple degradation of ER lumenal contents by vacuolar proteases is not the reason for the LegC7-mediated growth inhibition.

### LegC7 expression alters the redox state of the ER lumen

To determine whether LegC7 expression altered the general physiology of the ER through a potential ER:endosome fusion event, we employed a redox-sensitive, ER-targeted GFP (eroGFP) to observe the redox state of the ER lumen during LegC7 expression. EroGFP contains a pair of cysteine residues that form a disulfide bond under oxidizing conditions, which reorients the chromophore. This reorientation decreases excitation at a 488 nm, and increases excitation at a 405 nm; the ratio of fluorescence from these two maxima serves as a measurement for the redox state of the ER lumen in vivo[34]; addition of either DTT (reducing) or H_2_O_2_ (oxidizing) to cells expressing eroGFP changes this ratio, as expected (**Fig. S5**). To determine the effects of LegC7 expression on the ER redox state, cells (*n*=10^5^) were passed through a flow cytometer 6 hours post-induction of LegC7 expression, and fluorescence from excitation at 405 nm and 488 nm was measured. Fluorescence from cells without eroGFP (**Fig. 7A**) and heat-killed, eroGFP-expressing cells (**Fig. 7B**) was measured to define populations of low-fluorescence cells and dead eroGFP-expressing cells, respectively. These populations were removed from data sets for each other strain (**Fig. 7C-F**) to isolate the populations used to determine the ratio of fluorescence (Ex_405_/Ex_488_) as a measure of ER redox state (**Fig. 7G-J**).

**Figure 7.**
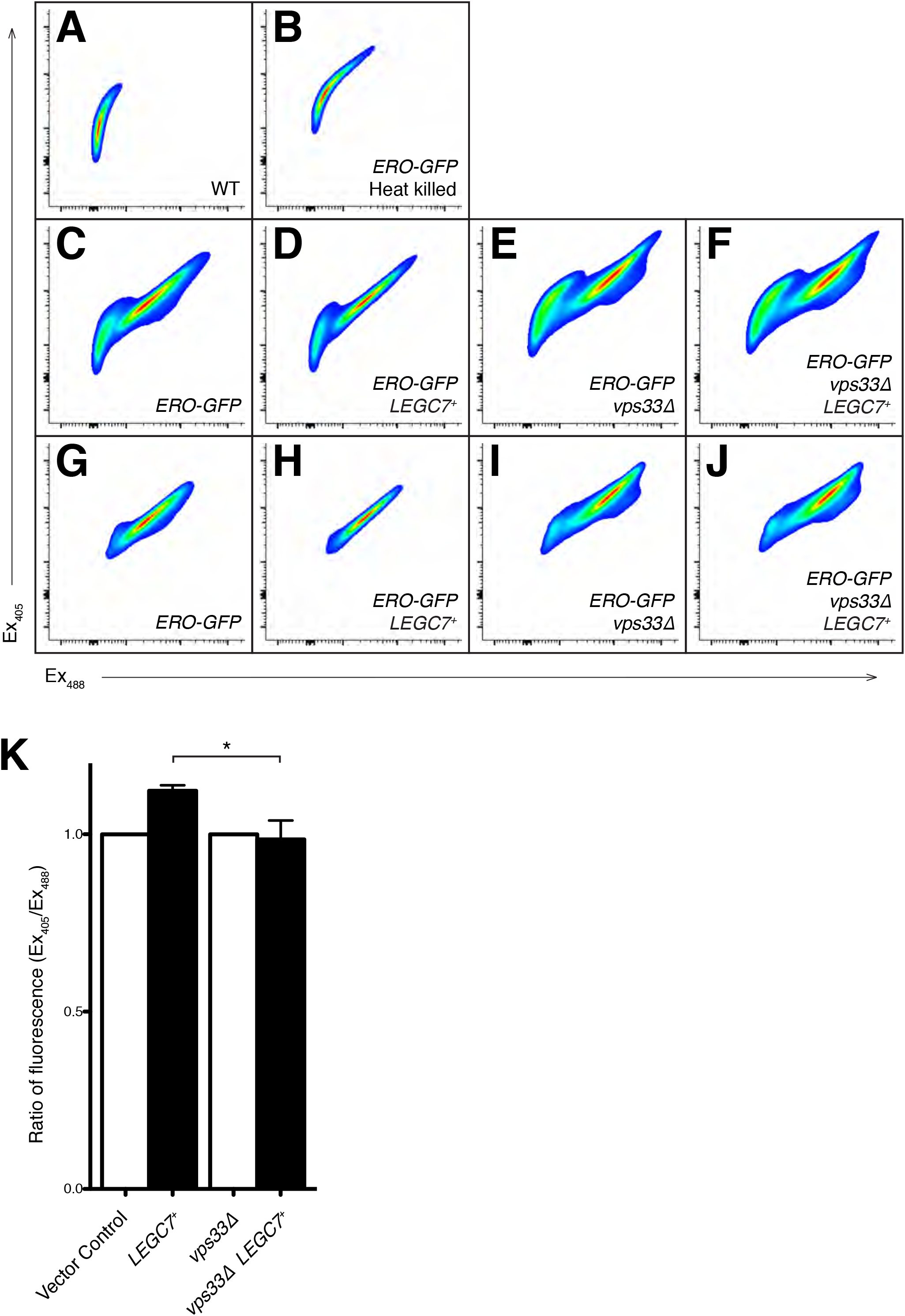
LegC7 expression oxidizes the ER lumen, but not in a *vps33*Δ mutant. Yeast strains containing ER-targeted redox-sensitive eroGFP were grown to saturation in CSM medium at 30°C, harvested via centrifugation, resuspended in an equal volume of fresh CSM containing 1% raffinose and 1% galactose, and grown for an additional 6 h at 30°C. Cells were then analyzed via flow cytometry (n = 10^5^). Gates for cells without eroGFP (**A**) and heat-killed cells containing eroGFP (**B**) were used to eliminate low fluorescence and dead cell populations, respectively, from strains in panels (**C** – **F**) to generate panels (**G** – **J**). (**K**) Ratios of GFP fluorescence (Ex_405_/Ex_488_) were calculated, and ratios for (**G**) and (**H**) were normalized by the same factor such that the ratio for (**G**) = 1. ratios for (**I**) and (**J**) were normalized such that the ratio for (**I**) = 1. Three independent experiments were performed to generate ratios displayed in (**K**); error bars represent ± the standard deviation between experiments; (*) = P < 0.05. Panels (**A** – **J**) are representative plots, taken from one experiment.

Typically, the ER maintains an oxidizing environment to facilitate the oxidative folding of nascent proteins[35]. After 6 hours of LegC7 expression, the ratio of fluorescence (Ex_405_/Ex_488_) increases ~10% compared to the vector control, corresponding to a higher percentage of oxidized eroGFP molecules (**Fig. 7K**), similar to the observed response to 10 mM H_2_O_2_ (**Fig. S5**). However, LegC7 expression in a *vps33*Δ strain does not change the ratio of fluorescence compared to the vector control (**Figs. 7I, J**, **and** **K**); suggesting that further oxidation of the ER lumen during LegC7 expression is dependent upon functional endosomal tethering complexes.

### LegC7 may induce ER:endosome fusion events

Based on our evidence suggesting LegC7 induces fusion of ER and endosomal compartments, we designed a split-GFP bimolecular complementation system to assess this possibility in vivo (**Fig. 8A**). It is known that the GFP chromophore can be divided into two separate, non-fluorescent domains (GFP_1-10_ and GFP_11_)[36]. However, upon being brought into close proximity to each other, the GFP_11_ helix can interact with GFP_1-10_, thereby reconstituting functional and fluorescent GFP[36]. Therefore, by directing these individual domains of GFP to the lumen of distinct membrane-bound compartments, we can measure reconstituted GFP fluorescence as a proxy for a fusion event. To measure the possibility of ER fusing to endosomal compartments, GFP_1-10_ was fused to the first 50 amino acids of the vacuolar carboxypeptidase Y protein (CPYss), which is known to be responsible for the targeting of CPY to the vacuole via endosomes[37]. In addition, the GFP11 helix was fused to the C-terminus of the transmembrane domain of integral ER membrane protein Scs2p (Scs2TM) to direct it to the lumenal side of the ER [36]. In order to confirm proper localization of these constructs, mRuby2 was fused to the N-terminus of Scs2TM-GFP_11_. CPYss-mRuby2-GFP_11_ was also constructed as a control to confirm the delivery of CPYss-GFP_1-10_ to endosomes and to control for endosomal compartment density during LegC7 expression (**Fig. 8A**).

**Figure 8.**
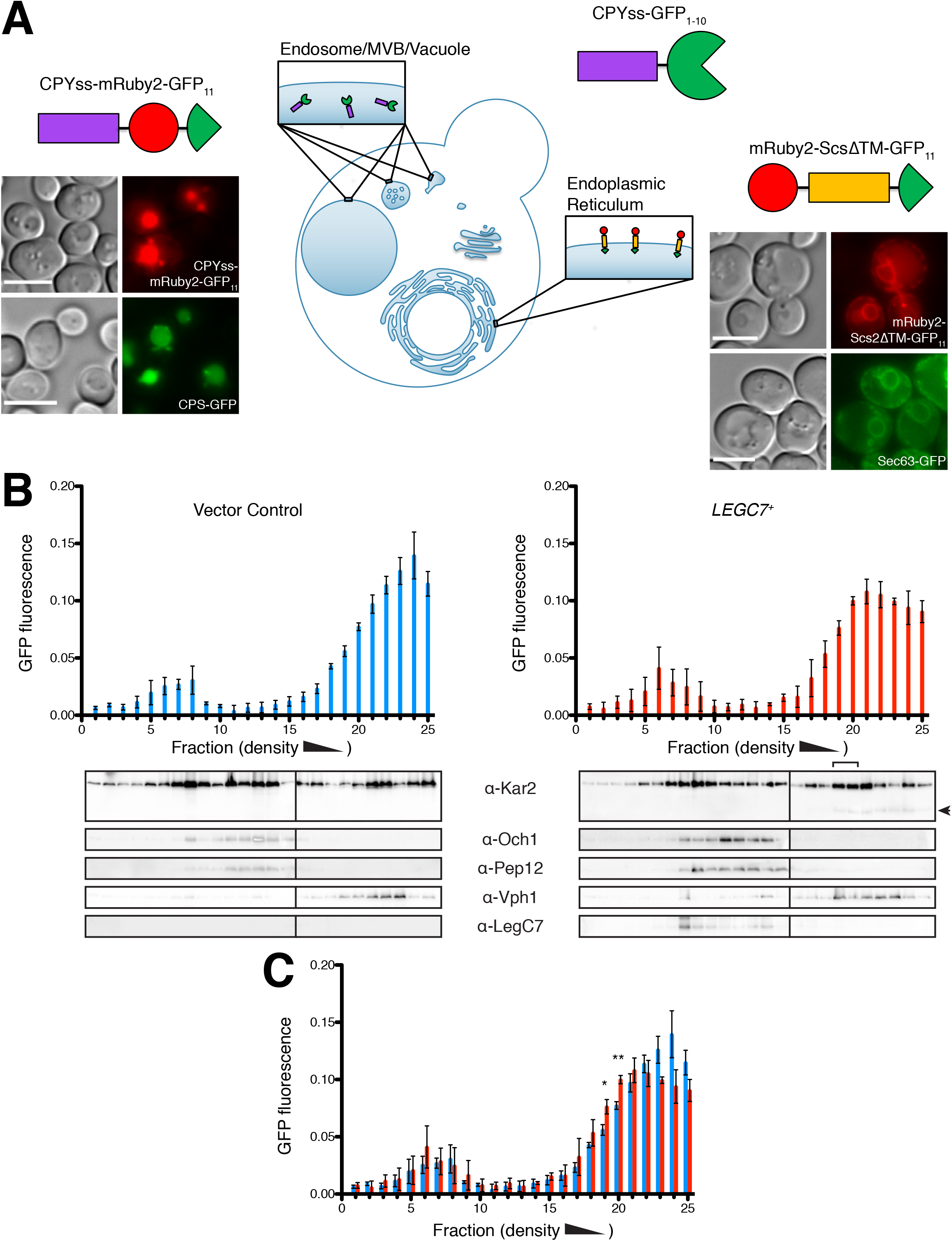
LegC7 may cause fusion of ER-derived and endosomal compartments. (**A**) Model of the split-GFP system constructed for this study. Strains containing individual constructs were grown to saturation in CSM medium at 30°C for imaging. Scale bars = 5 μ. (**B**) Strains containing ER-targeted mRuby2-ScsΔTM-GFP11 and endosome-targeted CPYss-GFP1-10 and either pYES2NT C (left) or pYES2-*LEGC7*^+^ (right) were grown to saturation in CSM medium at 30°C, harvested via centrifugation, resuspended in an equal volume of fresh CSM containing 1% raffinose and 1% galactose, and grown for an additional 18 h at 30°C. Equal amounts of each strain were fractionated into 25 fractions (Materials and Methods), and 20 μL volumes of each fraction were measured in triplicate for GFP fluorescence and averaged. Error bars represent ± the standard deviation across three independent experiments. The remainder of each fraction was TCA precipitated, and equal volumes were separated via SDS-PAGE and immunoblotted for the ER luminal ATPase Kar2p, the cis-Golgi mannosyltransferase Och1p, the early endosomal t-SNARE Pep12p, the vacuolar ATPase subunit Vph1p, and LegC7. The black arrow indicates Kar2 degradation product and the bracket indicates fractions 19 and 20 where we see Kar2p degradation and a shifted fluorescence peak. (**C**) GFP fluorescence plots were combined for comparison; (*) = P < 0.05; (**) = P < 0.005.

Cells harboring CPYss-GFP_1-10_ (endosomes), mRuby2-Scs2TM-GFP_11_ (ER), and a vector control were grown to mid-log, lysed, and membrane-bound compartments were subjected to density-dependent fractionation. Following fractionation, GFP fluorescence of each fraction collected was measured and plotted as a percentage of the total fluorescence across all fractions (**Fig. 8B**). The remainder of each fraction was TCA precipitated and probed for organelle markers and LegC7. In strains expressing LegC7, the maximum GFP fluorescence ratio peak shifts towards more dense compartments; this shift was highly reproducible across experiments (**Figs. 8B** **and** **C**). LegC7 expression had no effects on the fractionated GFP fluorescence from strains expressing CPYss-GFP_1-10_ and CPYss-mRuby2-GFP_11_ (both endosomal, **Fig. S6**). Interestingly, the two fractions with significantly higher GFP fluorescence during LegC7 expression, fractions 19 and 20 (**Fig. 8C**), were the densest fractions in which Kar2p degradation was first detected (**Fig. 8B**, **bracket**).

While LegC7 protein did not colocalize with compartments containing the Kar2p degradation product, LegC7 induced an apparent enrichment of early endosomal t-SNARE Pep12p in more dense fractions also containing ER membrane (Kar2p) and LegC7 (**Fig. 8B**, right panel), suggesting a shift of endosomal membranes to ER densities. Taken together, these results show that LegC7 expression causes the mixing of some ER and endosomal compartments, resulting in the reconstitution of a split-GFP protein and causing aberrant degradation of ER content by endosomal/vacuolar proteases.

## Discussion

To survive intracellularly, *L. pneumophila* translocates over 300 effector proteins into host cells via its Dot/Icm Type IVB secretion system, inducing dramatic changes to normal host ER and endolysosomal membrane trafficking pathways, such as forcing the aberrant activation of the host ER Rab1 GTPase [38, 39], inhibition of autophagy via RavZ[9], and inhibiting the function of the lysosomal V-type ATPase via the activity of SidK[40]. A critical component of LCV formation, however, is through the active recruitment of ER-derived membranes to the phagosomal membrane. While the precise mechanism by which *Legionella* induces ER:phagosome fusion is not completely known, the activities of *Legionella* LegC7/LegC2 have been shown to be important for optimal biogenesis of the LCV via ER membrane recruitment, although deletions of these genes do not result in overall intracellular proliferation defects[14, 17]. Accordingly, characterizing the in vivo activities of these proteins has proven difficult. Our previous discovery that LegC7 expression disrupts endosomal trafficking pathways in yeast[15], coupled with the discovery that LegC7 colocalizes with KDEL proteins[14] and forms a membrane fusion-competent, NSF/α-SNAP-resistant, SNARE-like complex with *Legionella* LegC2, LegC3, and host VAMP4 proteins[18], led us to explore the effects of LegC7 expression on potential ER and endosomal fusion events in vivo.

Supporting the previous discoveries that localize LegC7 to ER-containing structures in mammalian cells[7, 14], we describe drastic ER morphology changes upon expression of LegC7 in yeast (**Fig. 2**). Furthermore, we find that LegC7 immunoprecipitations enrich for Emp47p/Emp46p/Ssp120p complex subunits, suggesting an interaction between LegC7 and specific host cargo adapters for biosynthetic cargo leaving the ER. Interestingly, *emp46*Δ *emp47*Δ yeast strains appear to phenocopy the defective carboxypeptidase S vacuolar delivery phenotype of LegC7-expressing strains (**Fig. 1**), indicating that LegC7 may inhibit Emp47p/Emp46p/Ssp120p complex function, either directly or indirectly. However, the fact that these *EMP46/47* deletions minimally suppress LegC7-mediated growth inhibition contradicts the notion that LegC7 simply functions to inhibit Emp47p/Emp46p, but rather indicates that defects in ER:Golgi cargo delivery appear to result in the same trafficking defects observed in LegC7-expressing strains. Perhaps host cargo adapters play a role in targeting LegC7 to ER exit sites and ER-derived compartments containing cargo beneficial to *L. pneumophila* during infection. Indeed, Emp46p and Emp47p are known to function in glycoprotein secretion[21], with a function similar to mammalian ERGIC-53/LMAN1[41], and glycoprotein acquisition has been shown to be important for nutrition and evasion of host immune defenses by bacterial pathogens, including *L. pneumophila*[42–44]. While there is no documented role for ERGIC-53 in intracellular bacterial infection, this possibility should be investigated in future studies. Although LegC7 was not determined to be glycosylated by yeast in these studies (**Fig. S2**), it may interact with these proteins via structural motifs, or be glycosylated in mammalian expression systems. Further experimentation to show direct interactions between LegC7 and Emp47p/Emp46p/Ssp120p complexes in vitro would be required to characterize these interactions.

Previous research has shown that, while LegC7 appears to localize to ER structures[7, 14], expression of LegC7 in yeast also induced the formation of large punctate structures on the vacuole membrane and caused the mis-secretion of vacuole-bound CPY-invertase; these phenotypes were suggestive of defects in endosomal maturation or multivesicular body formation[7]. Our current findings that LegC7 expression induces the formation of Sec63-positive, Vps8p-positive compartments or structures (**Fig. 5A** **and** **B**) now help reconcile these discoveries and indicate that LegC7 activity – in the absence of other *Legionella* effectors – either recruits, or fuses, ER-derived vesicles to endosomes. This activity is also likely to disrupt ER:Golgi trafficking, as implied by defects in GFP-CPS trafficking in *emp46*Δ*emp47*Δ strains (**Fig. 1**). As mass disruptions in ER:Golgi traffic would be lethal to yeast cells[45], this is likely the reason for the toxic nature of LegC7/YlfA upon expression in yeast[14].

Strikingly, deletions of genes encoding for the core subunits of the conserved endolysosomal membrane tethering complexes CORVET and HOPS provide complete resistance to LegC7-mediated toxicity (**Fig. 3A**). We also find that deletion of the CORVET-specific *VPS8* gene provides more robust suppression of toxicity than the other complex-specific subunits. This suggests a more prominent role for the endosomal CORVET complex in LegC7 activity, further supported by our observation of colocalization of Vps8p and LegC7 in vivo (**Fig. 4A** **and** **F**). In addition to suppressing LegC7 toxicity, deletion of class C core complex subunit *VPS33* prevents LegC7-mediated alteration of ER morphology (**Fig. 2**) and the downstream effects such as Kar2p degradation (**Fig. 6**) and enhanced oxidation of the ER lumen (**Fig. 7**). This, coupled with the finding that LegC7-induced Kar2 degradation is also dependent upon the early endosomal t-SNARE, Pep12p, provides evidence that LegC7 induces aberrant SNARE-dependent fusion of ER-derived membrane and endosomes in vivo. While it is well established that small GTPases are primarily responsible for recruiting tethering complexes, such as CORVET and HOPS, to their target membranes[27, 46, 47], it is worth noting recent evidence that SNAREs, to which LegC7 has been compared, also play a role in targeting tethering complexes to membranes[48]. The apparent genetic interaction between LegC7 and class C tethering complexes raises the possibility that these complexes, which are conserved in higher eukaryotes[46], are also utilized by *L. pneumophila* for optimal LCV biogenesis.

As stronger evidence of ER:endosome fusion observed in these studies, we found that Kar2p degradation is dependent upon active vacuolar proteases, showing authentic delivery of ER to degradative compartments; this degradation was not found to be due to ER-phagy or general autophagic pathways (**Fig. 6B**). Furthermore, endosomes and lysosomes are oxidative compartments[49] and the ER lumen is further oxidized with LegC7 expression, which cannot be attributed to ER stress induced by LegC7 expression, as the ER redox state remains the same when expressing LegC7 in a *vps33*Δ mutant. Lastly, the split-GFP assay developed for this study provides further evidence of LegC7-induced mixing of lumenal contents of ER-derived and endosomal compartments.

*L. pneumophila* regulates the activity of some Dot/Icm effectors with other Dot/Icm effectors, termed “metaeffectors”, by inactivation or marking target effectors for proteolytic degradation[50–52]. Multiple studies have shown that *L. pneumophila* accomplishes temporal regulation during infection by delayed secretion of metaeffectors[50] or continuous secretion into the host cytosol, titrating out the target effector over time[51]. A recent study showed that LegC7-mediated growth inhibition in yeast is suppressed by co-expression of the Dot/Icm substrate MavE[52]. Perhaps the suppression of growth inhibition is an indication that MavE regulates LegC7 during infection through inactivation or degradation. It is known that LegC7 is part of the subset of Dot/Icm effectors first secreted into the host, many of which affect vesicular traffic[53, 54], and MavE likely functions to provide the proper spatiotemporal regulation for LegC7 activity during *Legionella* pathogenesis, although whether or not MavE expression suppresses the LegC7-dependent ER:endosome fusion events observed in yeast has not yet been explored. This work provides evidence that LegC7 is capable of redirecting host biosynthetic traffic in a manner that is independent of other Dot/Icm substrates, but reliant on conserved host endolysosomal fusion machinery. Our findings are consistent with the notion that this redirected cargo is directed to the LCV, as the LCV lipid and protein composition has been compared to that of trans-Golgi and early endosomes[55].

Despite the strong link of LegC7 to the activities of CORVET and endosomal SNARE proteins, we were surprised to note that we never detected any co-precipitating yeast SNARE proteins or multisubunit tethering factors in our multiple LegC7 immunoprecipitations. While these interactions may be sufficiently transient so that they would be undetectable in these pulldowns, it is important to note that it has been suggested that LegC2/3/7 proteins engage host R-SNARE proteins in an NSF/a-SNAP-resistant complex[18]. Therefore, these interacting proteins may only be observed when all three LegC proteins are co-expressed. Nevertheless, LegC7 has an in vivo activity distinct from either LegC2 or LegC3, suggesting differences in localization and protein:protein interactions, at least in yeast but also possibly in higher organisms.

Importantly, SNARE proteins generally do not form fusion-competent complexes promiscuously in vivo, only forming complexes with specific combinations of other SNARE proteins, and the absence of any one SNARE protein prevents complex formation[16]. Therefore, if LegC2/3/7 form a specific and cognate 3Q-SNARE complex, we might expect limited functionality of these LegC proteins in the absence of other complex members. As such, the presence of observable phenotypes, let alone distinct phenotypes, produced by individual LegC proteins in yeast is generally uncharacteristic of SNARE proteins that form a 3Q-SNARE complex. Similarly, the fact that proteins that specifically antagonize LegC3 (LupA) and LegC7 (MavE) have been evolutionarily maintained in *Legionella* suggests that LegC3 and LegC7 probably function independent of each other during infection[52]. Furthermore, LegC2 and LegC7 form homooligomeric complexes through interaction between cytosolic domains[17], which is also distinct from SNARE proteins. These findings suggest potential multifunctionality and/or promiscuity of LegC proteins, which could be cost-effective for *Legionella* during infection. As such, future studies utilizing heterologous co-expression of combinations of LegC proteins or comparing phenotypes of *legC* single, double, and triple knockout mutants, as well as host CORVET/HOPS disruptions, could better our understanding of LegC protein functionality during *Legionella* pathogenesis.

## Materials and Methods

### Yeast strains and plasmid constructions

All yeast strains used in this study were derivatives of *Saccharomyces cerevisiae* strain BY4742 (MATα *his3*Δ*1 leu2*Δ*0 lys2*Δ0). Where denoted, expression of *GAL1*-based promoters was initiated with the addition of 1% raffinose and 1% galactose as the carbon source, in place of 2% glucose.

To create HaloTag^®^ LegC7 lacking the transmembrane domain, we amplified LegC7 using the primers LegC7HaloF and LegC7HaloF (**Table S1**) utilizing pVJS52 [13] as a template. The resulting PCR product and pHis6-Halo (Promega Corporation) were digested with PvuI and NotI and ligated to create pVJS77 that was confirmed via sequencing (Georgia Genomics Facility, University of Georgia).

To construct the vectors for expressing the components of the split-GFP bimolecular fluorescence complementation assay, a combination of overlap PCR and homologous recombination techniques were used (**Table S1**). To generate plasmid pRS415-*KAR2*_pr_-mRuby2-G_3_AS-*SCS2*TM-PWG_3_SM-GFP_11_ for directing the GFP_11_ helix to the ER lumen, four separate PCR fragments were amplified: Fragment A was amplified from BY4742 genomic DNA with the primer pair RSG F1 and RSG R1, Fragment B was amplified from pFA6a-link-yomRuby2-SpHis5 [56] with primer pair RSG F2 and RSG R2, Fragment C was amplified from BY4742 genomic DNA with primer pair RSG F3 and RSG R3, and Fragment D was amplified from pSJ1321 [36] with primer pair RSG F4 and RSG R4; resultant amplicons were then used as templates for overlap PCR. Fragments A and B were allowed to self-prime, then amplified with primer pair RSG F1 and RSG R2 to create Fragment E. Fragments C and D were allowed to self-prime, then amplified with primer pair RSG F3 and RSG R4 to generate Fragment F. Finally, Fragments E and F were allowed to self-prime, then amplified with primer pair RSG F1 and RSG R4 to create the final amplicon. This final amplicon was co-transformed into BY4742 with pRS415 (*CEN/ARS, LEU2*^+^) which had been previously linearized with XhoI and SpeI, using standard lithium acetate techniques[57].

To generate plasmid pRS425-CPY_pr+1-50_-mRuby2-[G_3_S]_2_LE-GFP_11_ for directing the GFP_11_ helix to endosomes/lysosomes, three separate amplicons were generated: Fragment G was amplified from BY4742 genomic DNA with primer pair CRG F1 and CRG R1, Fragment H was amplified from pFA6a-link-yomRuby2-SpHis5 with primer pair CRG F2 and CRG R2, and Fragment I was amplified from pSJ1321 with primer pair CRG F3 and RSG R4. Resultant amplicons were all mixed, allowed to self-prime, and the final amplicon was amplified with primer pair CRG F1 and RSG R4. This final amplicon was co-transformed into BY4742 with pRS425 (2μ*, LEU2*^+^) which had been previously linearized with XhoI and SpeI.

To generate plasmid pRS423-CPY_pr+1-50_-GFP_1-10_ for directing the non-fluorescent GFP_1-10_ protein to endosomes/lysosomes, two separate amplicons were generated: Fragment J was amplified from BY4742 genomic DNA with primer pair CRG F1 and CG R1, and Fragment K was amplified from pSJ1726 [36] with primer pair CG F2 and CG R2. Resultant amplicons were mixed, allowed to self-prime, and the final amplicon was amplified with primer pair CRG F1 and CG R2. This final amplicon was co-transformed into BY4742 with pRS423 (2μ*, HIS3*^+^) which had been previously linearized with XhoI and SpeI. All plasmid constructs were confirmed by sequencing (Eton Bioscience, Inc.).

### Reagent preparation

To prepare the α-LegC7 resin used for immunoprecipitation, Rosetta 2 (DE3) pLysS cells (Novagen^®^) containing pVJS77 were grown in Terrific Broth (TB) to an OD of 1.5-2. Expression of LegC7ΔTM-Halo was induced by addition of 1mM IPTG, and cells were grown at 18° C for 18 hours. Cells were harvested by centrifugation and cell pellets were disrupted by a single pass through a One Shot Cell Disruptor (OS Model, Pressure BioSciences Inc.) at 20,000 psi. Lysates were cleared (20 min, 20,000 x *g*, 4°C) after the addition of 1 mM PMSF and DNase. The resulting supernatant was applied to 500 μL Halo resin and incubated for 1 hour at 4°C. The resin was washed with 30 mL of Halo buffer (50mM HEPES pH 6.5, 300 mM NaCl, and 0.5 mM EDTA) before a 30 min incubation at 22°C with Halo buffer containing 1 mM ATP and 10 mM MgCl_2_. The column was then washed with 30 mL of Halo buffer containing 1% Triton before a final wash of 30 mL Halo buffer. Polyclonal antiserum raised against LegC7[15] was then passed over the column and washed with 10 mL of IgG binding buffer (100mM Sodium Phosphate, 150 mM NaCl). Bound antibodies were then eluted using 1 mL of elution buffer (0.2 M Glycine pH 2.0), and then conjugated to IgG resin using the Pierce Protein A IgG Plus Orientation Kit (Thermo Fisher), according to manufacturer’s instructions. Bound antibodies were cross-linked using disuccinimidyl suberate (DSS) and any additional DSS sites were blocked with 0.1M ethanolamine.

### Cell lysis, subcellular fractionation, and split-GFP fluorescence assay

Strains harboring the *GAL*-inducible pYES2-NT C control vector or pYES2-*LEGC7*^+^ plasmid[13] were grown to saturation at 30°C in CSM-uracil medium. Cells were then harvested via centrifugation, resuspended in an equal volume of fresh CSM-uracil medium containing 1% raffinose and 1% galactose, and grown for an additional 18 h at 30°C. 50 OD_600_ units were harvested from each condition via centrifugation, then washed with 10 mL of azide buffer (50 mM Tris-HCl pH 7.5, 10 mM NaN3) to halt energy-dependent processes. Cells were resuspended in 5 mL of disulfide reduction buffer (100 mM Tris-HCl pH 9.4, 10 mM NaN3, 50 mM β-mercaptoethanol) and incubated at room temperature for 10 minutes. Cells were then washed with 3 mL spheroplasting buffer (1 M sorbitol, 40 mM HEPES-NaOH pH 7.5, 10 mM NaN_3_), harvested, then suspended again in 3 mL of spheroplasting buffer containing 100 μg oxalyticase (Zymolyase®-20T, AMSBIO). Mixtures were incubated at 37°C for 1 hr with gentle shaking, then harvested via centrifugation. Spheroplasts were then gently suspended in 3 mL of lysis buffer (200 mM sorbitol, 50 mM KOAc, 2 mM EDTA, 20 mM HEPES-NaOH pH 6.8) containing a protease inhibitor cocktail (Pierce™ Protease Inhibitor Tablet, EDTA-free, Thermo Fisher Scientific), and transferred to a Dounce homogenizer on ice. Spheroplasts were homogenized with a tightly-fitting pestle (15 strokes), and lysates were cleared by centrifugation (2 times, 5 min, 500 x *g* at 4°C) and collected.

2 mL of each lysate prepared above was applied to the top of a discontinuous Histodenz density gradient, prepared in lysis buffer (1 mL 43%, 1 mL 37%, 1 mL 31%, 1.5 mL 27%, 1.5 mL 23%, 1.5 mL 20%, 1 mL 17%, 1 mL 13%, 1 mL 8%). Samples were centrifuged in polyallomer tubes (175000 x *g*, 18 h, 4°C), then secured to a ring stand. The bottom of each tube was pierced with an 18 g needle, and the samples were removed with a peristaltic pump set to 1.5 ml min^−1^. Flow was directed to a fraction collector set to collect 500 μl fractions (Retriever^®^ 500, Teledyne ISCO), which were then saved on ice.

In order to detect GFP fluorescence resulting from the reconstitution of GFP from the non-fluorescent GFP_1-10_ and GFP_11_ fragments, 20 μl of each fraction collected above was loaded, in triplicate, to a black 384-well plate (Corning®) and fluorescence was measured (λ_ex_ = 479; λ_em_ = 520) on a plate-reading fluorimeter (SynergyMX, BioTek, Gen5 v. 2.09). Relative fluorescence values for each fraction are reported as the average across the three samples for each fraction measured.

### Western blotting

Total protein from cell lysates and subcellular fractions from indicated strains were precipitated with the addition of trichloroacetic acid (final 2% w/v) and placed on ice for 10 min. Protein was collected via centrifugation (10 min, 17000 x *g*, 4°C), washed twice with 300 μL ice-cold acetone, and allowed to dry at room temperature for 10 minutes. Dry pellets were solubilized in 2x SDS Dye (100 mM Tris-HCl pH 6.8, 200 mM DTT, 4% SDS, 0.2% bromophenol blue, 20% glycerol) and incubated at 98°C for 10 min. Equal volumes of samples were separated on 13% SDS-PAGE gels and immunoblotted for indicated proteins using polyclonal or monoclonal antibodies (1:1000) in blocking buffer (0.1% Tween-20 and 5% dry non-fat milk). Goat anti-rabbit (1:20000, Thermo Fisher Scientific) and anti-mouse (1:20000, Thermo Fisher Scientific) IgGs conjugated to HRP were used as secondary antibodies. Blots were developed with SuperSignal™ West Pico PLUS chemiluminescent substrate (Thermo Scientific).

### Microscopy

Strains were grown to saturation at 30°C in selective media. Cells were harvested and resuspended in fresh media with or without inducer (1% raffinose and 1% galactose, where indicated) and grown for an additional 18 hr at 30°C. Strains were harvested via centrifugation, washed, then mounted to glass slides which had been pretreated with a 1:1 mixture of concanavalin A (2 mg ml^−1^):polylysine (10% w/v). Slides were imaged with a Nikon Ti-U fluorescence microscope, equipped with appropriate filter sets. Images were processed with the Fiji software package[58, 59].

### Flow Cytometry

To measure the redox environment of the ER-localized eroGFP protein, indicated yeast strains were transformed with pPM28[34], and resultant strains were grown to saturation at 30°C in selective media. Cells were harvested, resuspended in fresh media with inducer (1% raffinose and 1% galactose), and grown for an additional 6 hrs at 30°C. 100,000 cells per strain were analyzed with a BD Biosciences LSR II flow cytometer (UGA Flow Cytometry Core), exciting with 405 nm and 488 nm lasers, and collecting fluorescence between 500-550 nm. Cells treated with DTT were analyzed 20 minutes after addition of DTT to the media, and cells treated with hydrogen peroxide were analyzed 5 minutes after addition of hydrogen peroxide to the media. Data analysis was performed within FlowJo 10.6. Gates defining populations of low fluorescence cells and dead cells were created with the Autogating tool, adjusted to maximum size (>98% of cells).

### Statistical Analysis

Statistical analysis was performed within the Prism software package (GraphPad Software, version 5.0f). Column statistics were performed via one-tailed, unpaired t-tests with Welch’s correction.

## Acknowledgments

The authors would like to thank Dr. Alexey Merz for providing essential reagents and Drs. Greg Odorizzi and Youngsoo Jun for providing essential insight and suggestions. pFA6a-link-yomRuby2-SpHis5 was a gift from Wendell Lim and Kurt Thorn (Addgene plasmid # 44858). pSJ1321 (pRS315-NOP1pr-GFP11-mCherry-PUS1) and pSJ1726 (pFA6-NATMX-CDC42pr-KAR2SS-GFP1-10) were gifts from Sue Jaspersen (Addgene plasmids # 86413 and # 86421). YCPlac33-sGFP-VRG4 was a gift from Benjamin Glick (Addgene plasmid # 25447). V.J.S. was supported by a grant from the National Institute of Allergy and Infectious Diseases (R01-AI100913). This research was supported in part by the National Institutes of Health grants 1S10OD018530 and P41GM10349010 to the Complex Carbohydrate Research Center. The Authors have no conflicts of interest to declare.

