## Supplemental Materials for "*Legionella pneumophila* LegC7 effector protein drives aberrant ER:endosome fusion in yeast"

**Methods:**

**Protease digestion and LC-MS/MS analysis of Leg C7 sample.**

**Precipitation of protein from SDS buffer and reduction/carbamidomethylation:** Cold acetone was added to the protein mixture in SDS buffer and incubated overnight at -20 °C. The precipitate obtained was recovered by centrifugation and washed with cold acetone. Protein precipitate was redissolved in digestion buffer (50 mM aq. $NH_4CO_3$), reduced by DTT (25 mM for 45 min), carbamidomethylated by Iodoacetamide (90 mM for 45 min) and dialyzed against dd$H_2O$.

**Trypsin digestion:** 25 µL of digestion buffer (50 mM aq. $NH_4CO_3$) was added to 20 µL of Sample (~ 20.0 µg) protein solution. The protein sample was digested by adding 5 µL sequencing-grade trypsin (Promega, 0.5 µg/µL) and incubated at 37 ºC for 24 h. The digests were desalted by C18 centrifuge cartridges. The digests in elution buffer (80 % acetonitrile and 0.1 % formic acid) were dried under speed vac. The peptides and glycopeptides were subsequently re-dissolved in solvent A (0.1% formic acid in water) and stored at - 30 ºC until analysis by nano-LC-MS/MS.

**Data acquisition of protein digest samples using nano-LC-MS/MS:** Desalted peptides were analyzed on an Orbitrap Fusion instrument (Thermo Scientific) equipped with a nanospray ion source with CID, HCD and ETD fragmentation options and connected to a Dionex binary solvent system. Pre-packed nano-LC columns of 15 cm length with 75 µm internal diameter (id), filled with 2 µm C18 material (reverse phase) were used for chromatographic separation of samples. After the precursor ion scan at 120000 resolution in Orbitrap analyzer, precursors at a time frame of 3 sec were selected for subsequent fragmentation using HCD at normalized collision energy of 28. Another acquisition with a program HCD product triggered ETD, where ETD fragmentation occurs based on the presence of glycan oxonium ions in the HCD fragmentation of the same peptide, was also employed. The threshold for triggering an MS/MS event on ion-trap was set to 500 counts. Charge state screening was enabled, and precursors with unknown charge state or a charge state of +1 were excluded (positive ion mode). Dynamic exclusion was enabled (exclusion duration of 60 s). The fragment ions were analyzed on orbitrap for HCD at 30000 resolution.

**Peptides and glycopeptide analysis:** The .raw files of the LC-MS/MS acquisition were analyzed through Byonic v2.6 and Proteome Discoverer 1.4 software against the .fasta sequence of LegC7 sample. Search was conducted with modifications such as oxidation of methionine, carbamidomethylation of cysteine and possible *N*-glycans from the corresponding expression species. A precursor ion tolerance of 10 ppm and fragment ion tolerance of 0.1 Da was set for the search with up to two missed cleavage for the target enzyme trypsin. Based on the identifications of the software and manual validation of spectra, sequences of amino acids on the peptides were validated. The HCD $MS^2$ spectra of glycopeptides were evaluated for the glycan neutral loss pattern, oxonium ions and the glycopeptide fragmentations to assign the sequence and the presence of glycans in the glycopeptides.

**Table S1. Primers used in this study[a].**

| | | |
|---|---|---|
| RSG F1 | 5’-CGACTCACTATAGGGCGAATTGGGTACCGGGCCCCCCCTCGAGCAGCCACCAGCCGC | pRS415-*KAR2* promoter-mRuby2 |
| RSG R1 | 5’-GGACACCATGGTATGTTTGATACGCTTTTTCCC | |
| RSG F2 | 5’-GCGTATCAAACATACCATGGTGTCCAAAGGAGAGG | mRuby2-$G_3$AS |
| RSG R2 | 5’-GCTAGCACCACCACCCTTATACAATTCATCCATACCAC | |
| RSG F3 | 5’-GGTGGTGGTGCTAGCTCATCCAGCATGGGTATATTC | $G_3$AS-*SCS2*TM-PW$G_3$SM |
| RSG R3 | 5’-CATAGAACCACCACCCCATGGTCTGTAGAACCATCCTAAAACC | |
| RSG F4 | 5’-CCATGGGGTGGTGGTTCTATGAGAGATCATATGGTTTTGCATG | PW$G_3$SM-$GFP_{11}$-pRS415 |
| RSG R4 | 5’-GAGCTCCACCGCGGTGGCGGCCGCTCTAGAACTAGTTTAAGTAATACCAGCAGCATTAAC | |
| CRG F1 | 5’-CTCACTATAGGGCGAATTGGGTACCGGGCCCCCCCTCGAGCATAGCGATGTTGGTCATCC | pRS425-CPY promoter + ss-mRuby2 |
| CRG R1 | 5’-GGACACCATGGAGAGATGATCCAGGTCGAG | |
| CRG F2 | 5’-GGATCATCTCTCCATGGTGTCCAAAGGAGAGGAG | mRuby2-$[G_3S]_2$LE |
| CRG R2 | 5’-CCATATGATCTCTCTCGAGAGAACCACCACCAGAACCACCACCCTTATACAATTCATCCATACCACC | |
| CRG F3 | 5’-GTATAAGGGTGGTGGTTCTGGTGGTGGTTCTCTCGAGAGAGATCATATGGTTTTGCATG | $[G_3S]_2$LE-$GFP_{11}$-pRS425 (with RSG R4) |
| CG R1 | 5’-CACCTTTAGACATGGAGAGATGATCCAGGTCGAG | pRS423-CPY promoter- + ss $GFP_{11}$ (with CRG F1) |
| CG F2 | 5’-CTCGACCTGGATCATCTCTCCATGTCTAAAGGTGAAGAATTGTTTAC | $GFP_{11}$-pRS425 |
| CG R2 | 5’-GAGCTCCACCGCGGTGGCGGCCGCTCTAGAACTAGTTCATCGATGAGAACCACC | |

[a]$G_3$As, PW$G_3$SM, and $[G_3S]_2$LE are single-letter amino acid codes for linker sequences added. ss = first 50 amino acids encoding the signal sequence of CPY (*PRC1*).

**Table S2. Complete protein ID list from LegC7 immunoprecipitations.**

| Protein Name | Score | # of Peptides |
|---|---|---|
| Emp47 | 3050.32 | 32 |
| Emp46 | 1277.74 | 22 |
| Sro9 | 923.29 | 9 |
| Atp2 | 821.02 | 9 |
| Glycine tRNA ligase | 800.08 | 13 |
| Hsc82 | 737.75 | 8 |
| Ils1 | 666.45 | 13 |
| Ssp120 | 481.79 | 7 |
| Idh2 | 474.07 | 6 |
| Mir1 | 403.57 | 6 |
| Crn1 | 389.80 | 9 |
| Ilv2 | 381.30 | 6 |
| Adt2 | 362.39 | 5 |
| YHR020W | 317.44 | 7 |
| Hrk1 | 298.80 | 8 |
| Tub2 | 270.01 | 4 |
| Rna1 | 223.91 | 5 |
| Sam1 | 207.93 | 2 |
| Pma1 | 194.12 | 4 |
| Srp40 | 190.52 | 4 |
| Gus1 | 174.56 | 2 |
| Rpl3 | 166.36 | 2 |
| Boi1 | 148.85 | 3 |
| Atp1 | 148.04 | 1 |
| Lys12 | 134.47 | 3 |
| Faa1 | 132.03 | 2 |
| Rps20 | 119.97 | 2 |
| Fas1 | 113.02 | 1 |
| Cpn60 (*Legionella*) | 109.51 | 2 |
| Gpp1 | 108.84 | 1 |
| Tub1 | 100.28 | 2 |
| Dbp3 | 88.61 | 2 |
| Gfa1 | 79.46 | 2 |
| Kap123 | 78.26 | 1 |
| Nsr1 | 75.33 | 1 |
| Rps0A | 74.93 | 1 |
| Ade5,7 | 73.31 | 2 |
| Nop58 | 70.61 | 1 |
| Rps11B | 62.67 | 1 |
| Hfa1 | 62.53 | 1 |
| Sdd3 | 55.40 | 2 |
| Lp12_1736 (*Legionella*) | 53.54 | 1 |
| Rpl21B | 48.80 | 1 |
| Ggc1 | 44.62 | 1 |
| Cps1 | 44.59 | 2 |
| Tef4 | 43.64 | 1 |

**Table S3. Predicted tryptic peptide sequences of LegC7.**

| Position of cleavage site | Amino acid sequence[a] | No of Amino acid residues | Supplementary figure # |
|---|---|---|---|
| 17 | ~~M~~ATNETELQVLIQHDSK | 17 | A |
| 33 | STITTSSLDSTDKDPK | 16 | B |
| 54 | ASTGTPEIESASLAQIVDTQK | 21 | C |
| 60 | QLSQVK | 6 | |
| 79 | ESLESIVDSIAENPSLITR | 19 | D |
| 92 | AASAWGELPMWQK | 13 | E |
| 145 | VTGGLVLTAPTLAVGLFAHIGVLLVIGGVTGLTYTAGAIVLDDHHT CNVNIAK | 53 | |
| 169 | EGLFGLADLLQITIEALDAIR | 21 | F |
| 178 | FAEEIEK | 7 | G |
| 185 | NENLR | 5 | |
| 192 | LTDNIDR | 7 | H |
| 212 | LGNEVESLSAQVELYMEIEK | 20 | I |
| 226 | DTNEMEQTVK | 10 | J |
| 234 | LQESTTK | 7 | |
| 241 | QTDLLEK | 7 | K |
| 263 | SQLQLAEK | 8 | L |
| 271 | IAELHEVR | 8 | M |
| 280 | LSLGLEVQK | 9 | N |
| 306 | TLEGTVQTLTGTVIADEEQR | 20 | O |
| 311 | VSFQK | 5 | |
| 320 | KLDGFLNDK | 8 | P |
| 330 | QLSFDQVAER | 10 | Q |
| 339 | AEEELK | 6 | |
| 351 | QSNDR | 5 | |
| 357 | YSELLK | 6 | |
| 365 | QEQQVER | 7 | |
| 373 | LGLHK | 5 | |
| 404 | ENVKPSNDPVHSGLLSHGIYSTPK | 24 | |
| 411 | VTQPK | 5 | |
| 418 | VEVVEDR | 7 | R |
| 425 | QTIALVN | 7 | |

[a]LegC7 Peptides detected via LC-MS/MS highlighted in green; underlined sequences are predicted possible *N*-glycosylation sites (http://www.cbs.dtu.dk/services/NetNGlyc/)

#### Figure Legends

**Figure S1. Identification of LegC7 peptides by LC-MS/MS.** Amino acid sequence coverage obtained by the LC-MS/MS analysis of tryptic digest of LegC7 (highlighted in green). Sites of *N*-glycosylation predicted by Net-N-glyc web tool are underscored and no evidence for *N*-glycosylation was observed. Based on the LC-MS/MS data amino acid at the first position (M – Methionine) either might not be present in the protein sample or had undergone non-specific cleavage.

**Figure S2. HCD MS2 spectra from tryptic digest of LegC7**. Panel designation refers to peptides annotated in Table S3 (green).

**Figure S3. *EMP46/47* deletions do not significantly reduce LegC7-mediated growth inhibition.** Yeast deletion mutants harboring either pYES2NT C or pYES2-*LEGC7*$^{+}$ were grown to saturation in CSM at 30°C. For each strain, 1 $OD_{600}$ unit was harvested via centrifugation, resuspended in 1 mL of 0.9% NaCl, and serially diluted 1:10 four times. 5 µL of each dilution was plated onto CSM containing 2% glucose and CSM containing 1% galactose and 1% raffinose to induce LegC7 expression.

**Figure S4. LegC7-mRuby2 inhibits yeast growth to a degree comparable to LegC7.** (**A**) Yeast strains harboring LegC7 constructs in pYES2NT C were grown to saturation in CSM at 30°C. For each strain, 1 $OD_{600}$ unit was harvested via centrifugation, resuspended in 1 mL of 0.9% NaCl, and serially diluted 1:10 four times. 5 µL of each dilution was plated onto CSM containing 2% glucose and CSM containing 2% galactose to induce LegC7 expression. (**B**) The same strains were grown to saturation in CSM medium at 30°C, harvested via centrifugation, resuspended in an equal volume of fresh CSM containing 2% galactose, and grown for an additional 18 h at 30°C. Equal amounts of each strain were harvested, protein was extracted, and equal volumes of each extract were separated via SDS-PAGE and immunoblotted for LegC7. Asterisk identifies an anti-LegC7 cross-reactive protein from yeast lysates that serves as a loading control.

**Figure S5. EroGFP can be utilized as a redox sensor of the ER lumen.** Yeast strains containing ER-targeted, redox-sensitive eroGFP were grown to saturation, treated with varying concentrations of DTT or $H_2O_2$ and cells were analyzed via flow cytometry ($n = 10^5$). After low-fluorescence and dead cell populations were removed from the data sets (See **Fig. 7**), ratios of GFP fluorescence were calculated and normalized by the same factor such that the ratio for untreated cells = 1. Error bars represent ± the standard deviation across 3 independent experiments.

**Figure S6. LegC7 expression in yeast strains containing endosome-directed CPYss-$GFP_{1\text{-}10}$ and CPYss-mRuby2-$GFP_{11}$ does not affect GFP fluorescence of subcellular fractions.** Strains containing endosome-targeted CPYss-mRuby2-$GFP_{11}$ and endosome-targeted CPYss-$GFP_{1\text{-}10}$ and either pYES2NT C or pYES2-*LEGC7*$^{+}$ were grown to saturation in CSM medium at 30°C, harvested via centrifugation, resuspended in an equal volume of fresh CSM containing 1% raffinose and 1% galactose, and grown for an additional 18 h at 30°C. Equal amounts of each strain were dounced and fractionated into 25 fractions (Materials and Methods), and 20 μL volumes of each fraction were measured in triplicate for GFP fluorescence and averaged. Error bars represent ± the standard deviation across three independent experiments.

**Fig. S1**

~~M~~AT**NET**ELQVLIQHDSKSTITTSSLDSTDKDPKASTGTPEIESASLAQIVDTQKQLSQVKESLE
SIVDSIAE**NPS**LITRAASAWGELPMWQKVTGGLVLTAPTLAVGLFAHIGVLLVIGGVTGLTYTA
GAIVLDDHHTCNVNIAKRLKEGLFGLADLLQITIEALDAIRKKFAEEIEKFKNENLRLTDNIDR
LGNEVESLSAQVELYMEIEKMLRKDTNEMEQTVKKLQESTTKQTDLLEKNQKELSKIRKEYEKS
QLQLAEKIAELHEVRLSLGLEVQKAKTVAKTLEGTVQTLTGTVIADEEQRVSFQKKLDGFLNDK
QLSFDQVAERICKAEEELKKVKEELRQSNDRYSELLKRQEQQVERLEKLGLHKLERIVDKENVK
PSNDPVHSGLLSHGIYSTPKGKVTQPKVEVVEDRQTIALVN

#### Figure S2

**A**

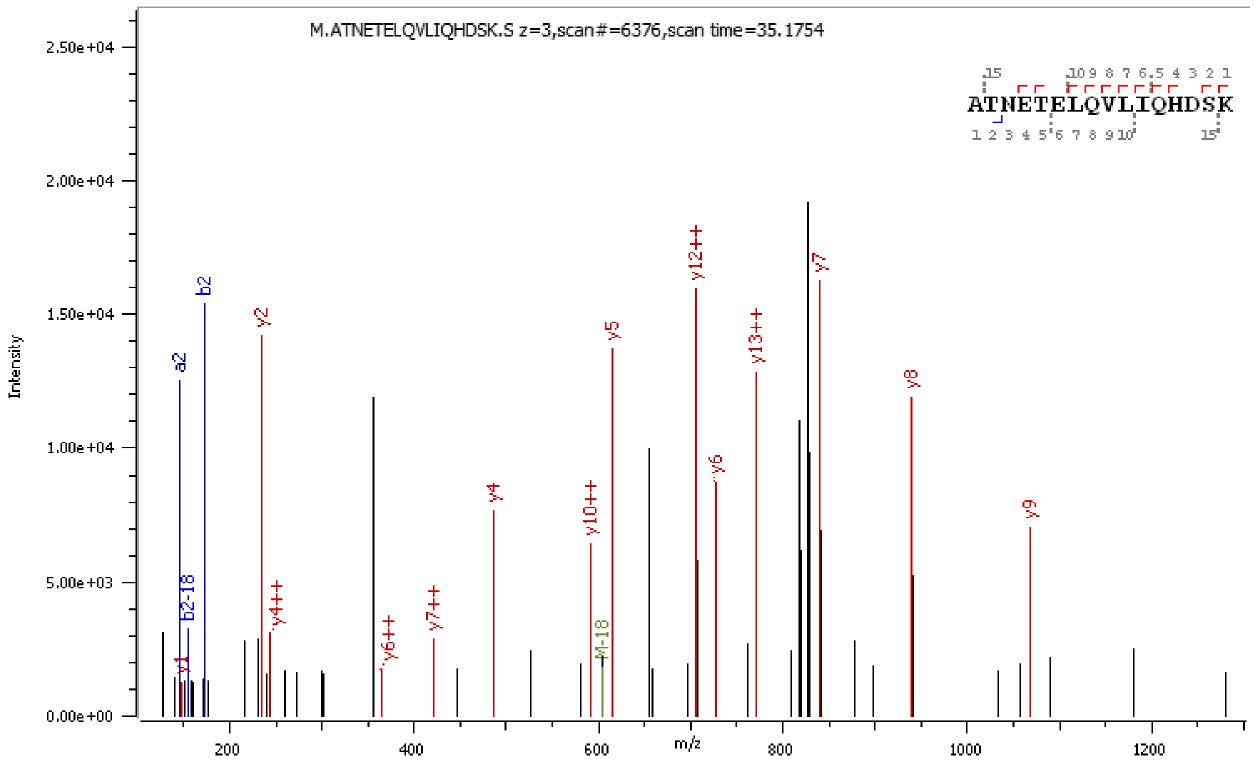

M.ATNETELQVLIQHDSK.S z=3,scan#=6376,scan time=35.1754
ATNETELQVLIQHDSK
Intensity
m/z


**B**

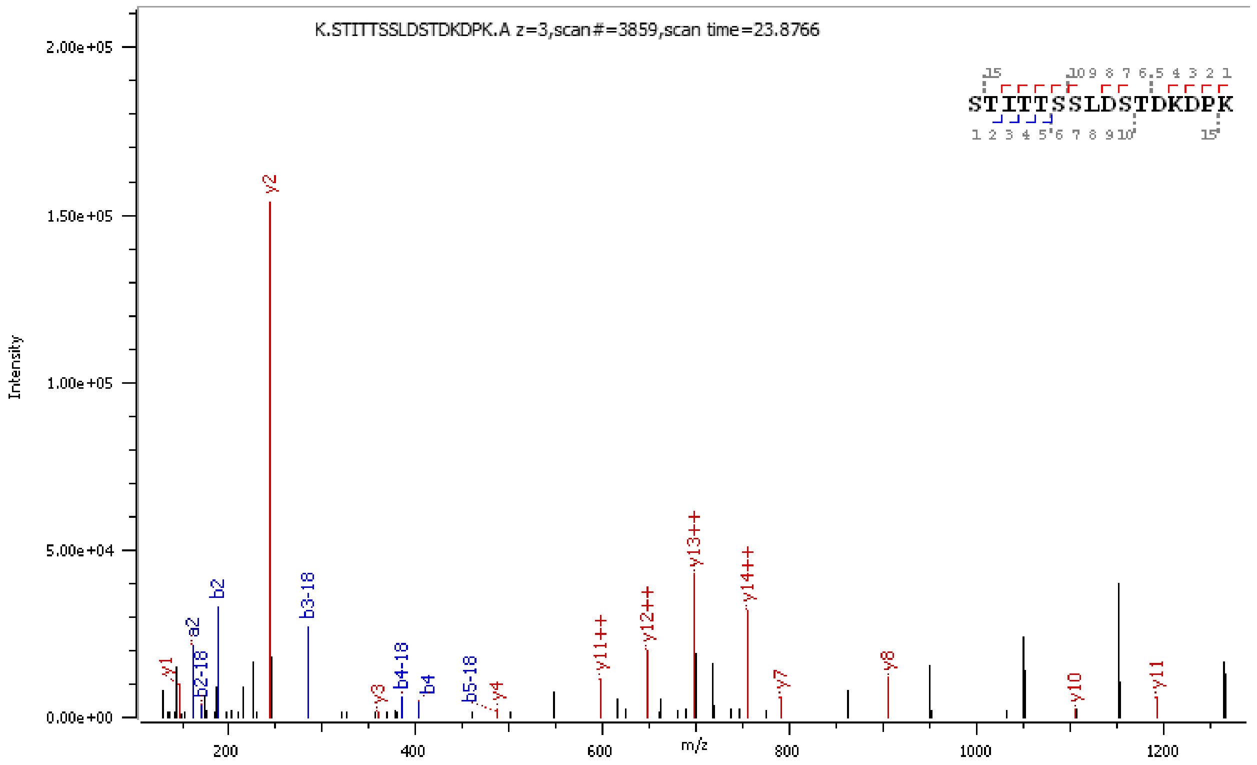

K.STITTSSLDSTDKDPK.A z=3,scan#=3859,scan time=23.8766
STITTSSLDSTDKDPK
Intensity
m/z

**C**

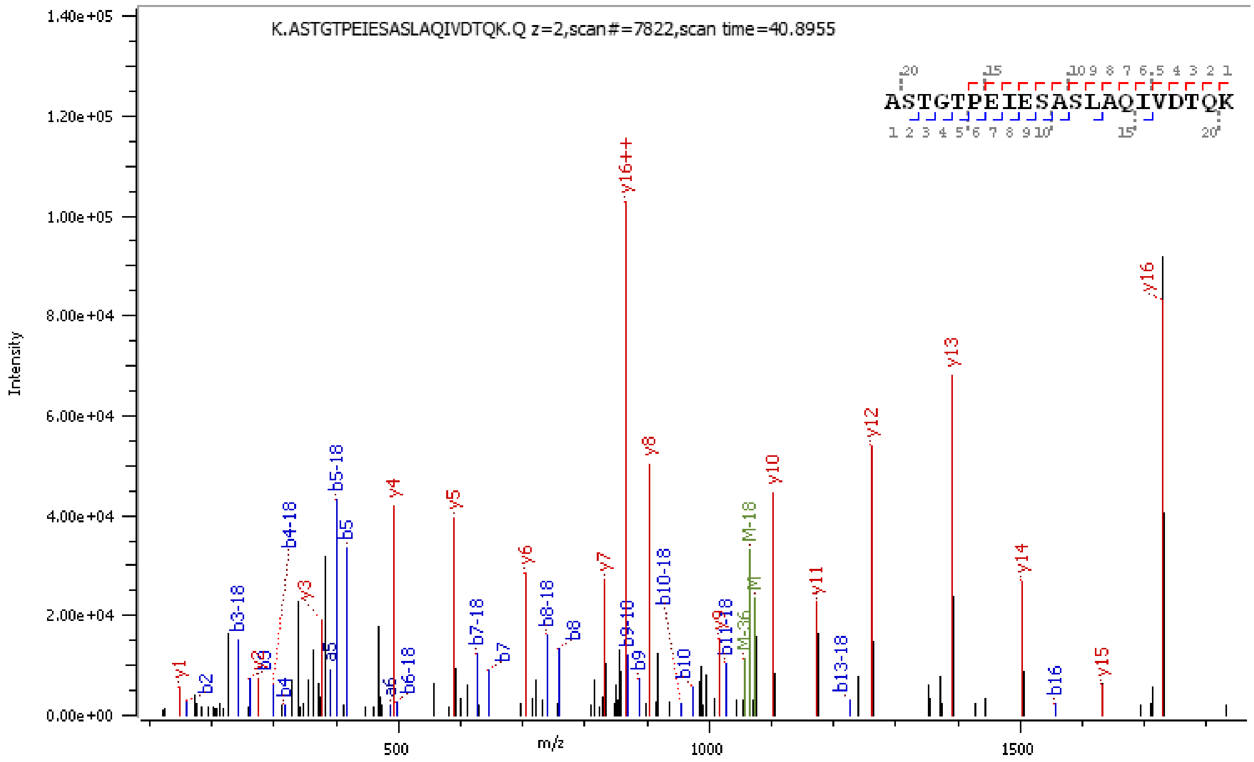
K.ASTGTPEIESASLAQIVDTQK.Q z=2,scan#=7822,scan time=40.8955
ASTGTPEIESASLAQIVDTQK
Intensity
m/z
1.40e+05
1.20e+05
1.00e+05
8.00e+04
6.00e+04
4.00e+04
2.00e+04
0.00e+00
500
1000
1500
y16++
y16
y13
y12
y8
y10
y5
y4
b5-18
b5
b4-18
y6
y7
b10-18
M-18
y11
y14
y15
b16
b13-18

**D**

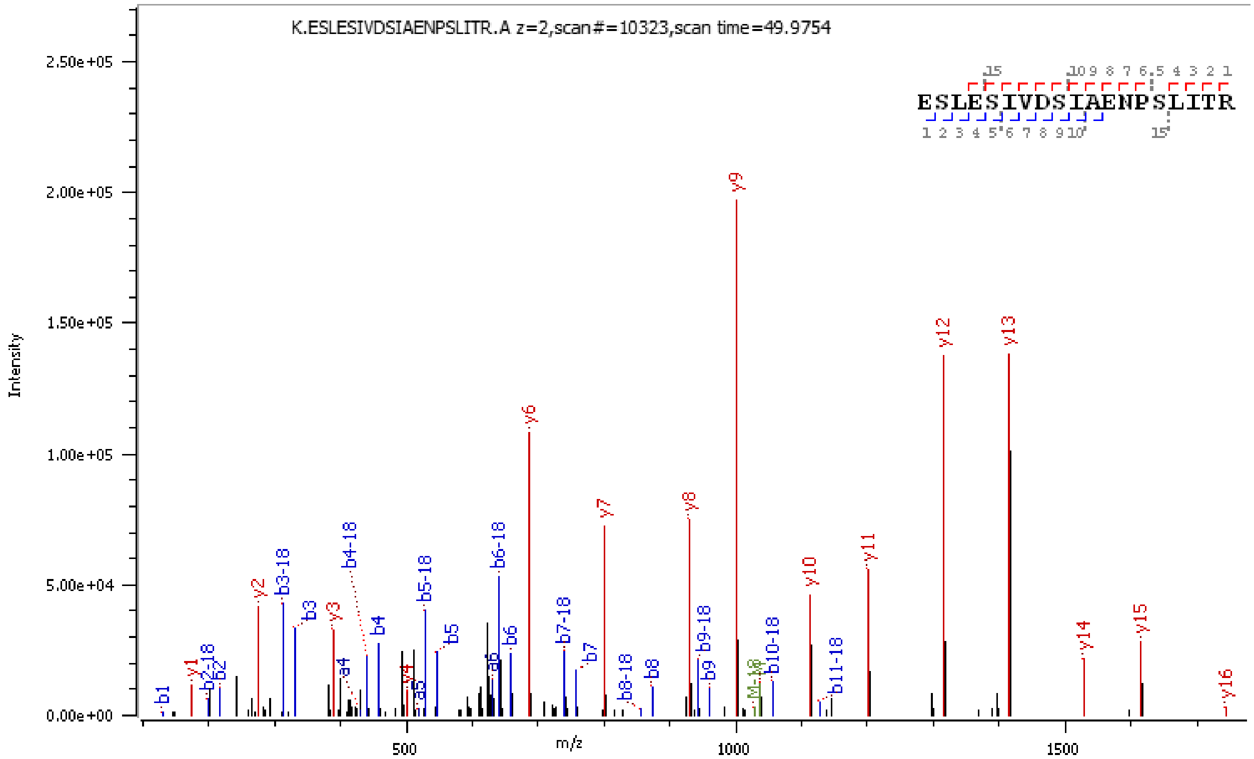
K.ESLESIVDSIAENPSLITR.A z=2,scan#=10323,scan time=49.9754
ESLESIVDSIAENPSLITR
Intensity
m/z
2.50e+05
2.00e+05
1.50e+05
1.00e+05
5.00e+04
0.00e+00
500
1000
1500
y9
y12
y13
y6
y7
y8
y10
y11
y14
y15
y16
y2
b3-18
b4-18
b5-18
b6-18
b7-18
b9-18
b10-18
b11-18

**E**

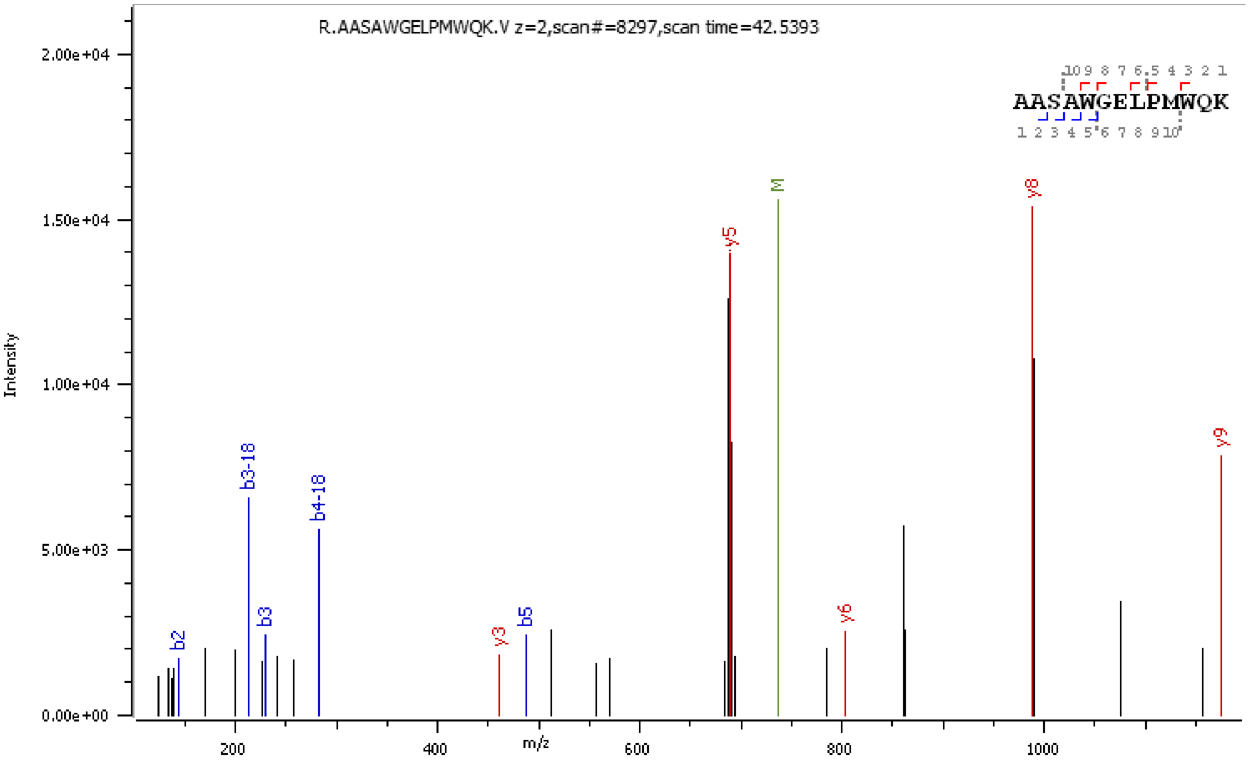
R.AASAWGELPMWQK.V z=2,scan#=8297,scan time=42.5393
AASAWGELPMWQK
Intensity
m/z
y5
M
y8
y9
b2
b3-18
b3
b4-18
y3
b5
y6

**F**

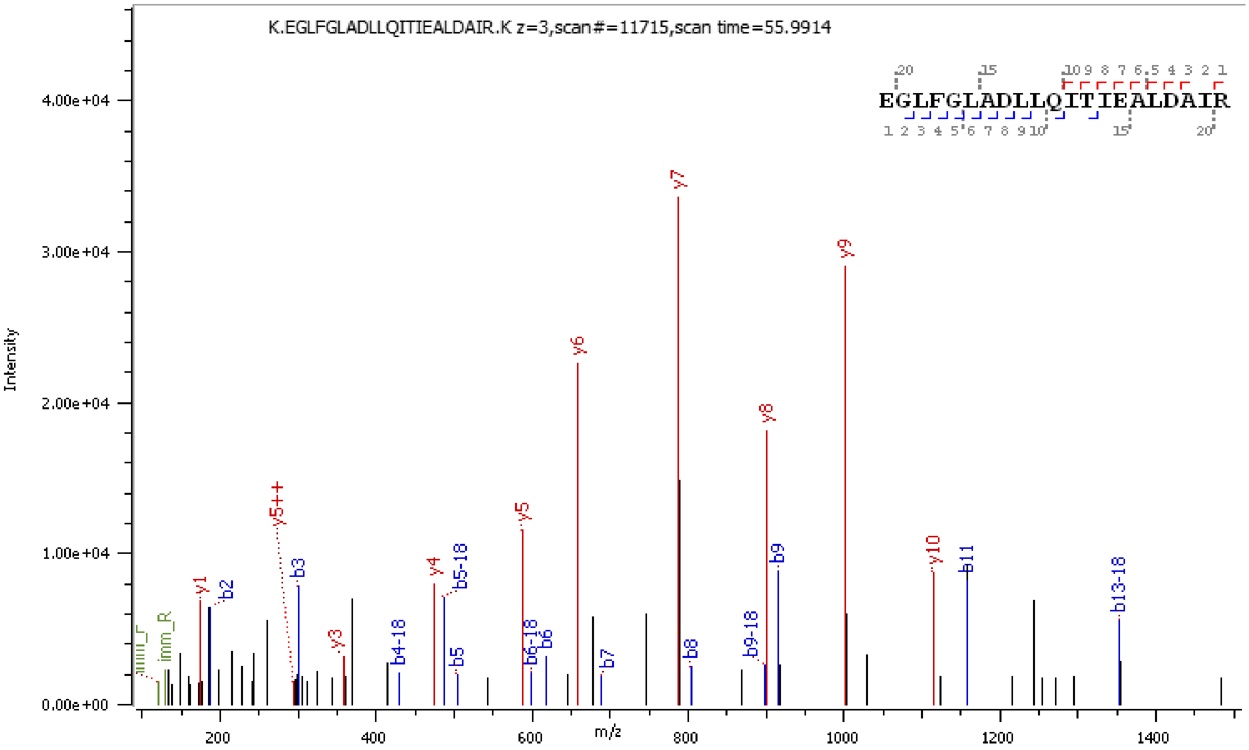
K.EGLFGLADLLQITIEALDAIR.K z=3,scan#=11715,scan time=55.9914
EGLFGLADLLQITIEALDAIR
Intensity
m/z
y7
y9
y6
y8
y5
y5++
y4
b5-18
b3
y1
b2
imm_R
y3
b4-18
b5
b6-18
b6
b7
b8
b9-18
b9
y10
b11
b13-18

G

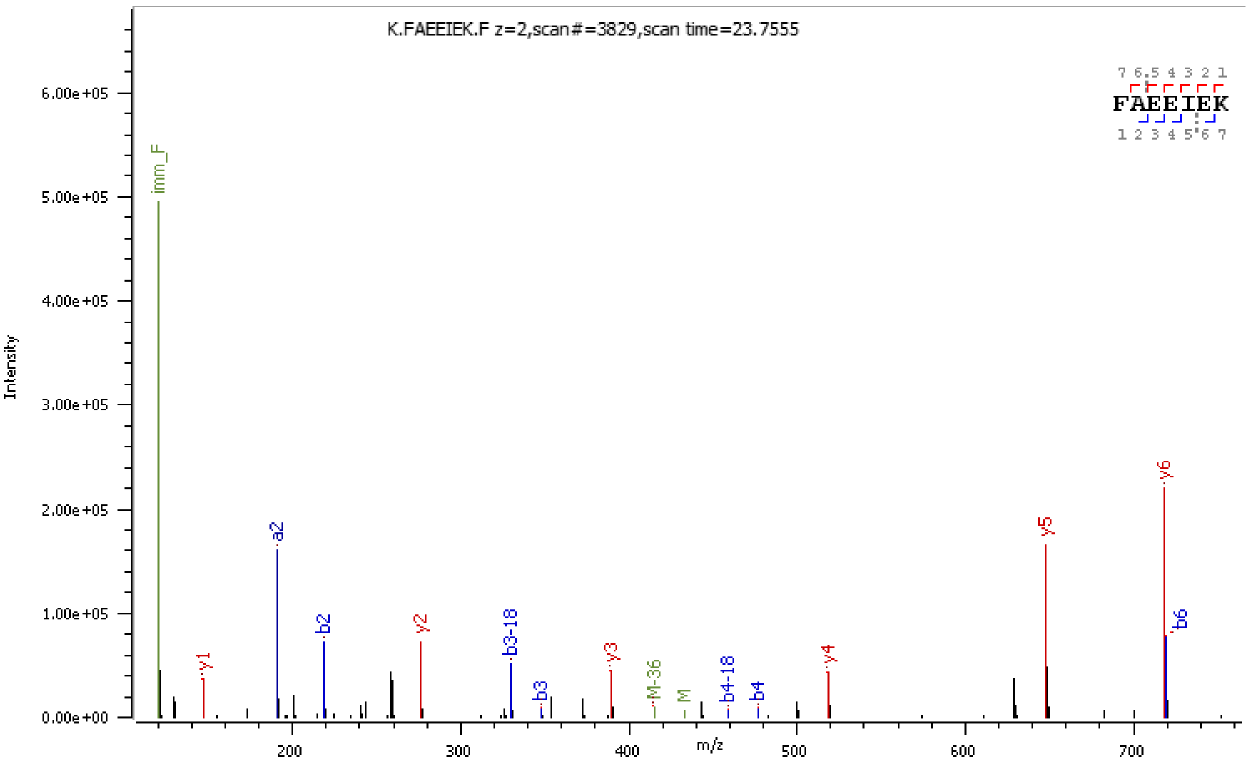
K.FAEEIEK.F z=2,scan#=3829,scan time=23.7555
FAEEIEK
Intensity
m/z
imm_F
a2
b2
y1
y2
b3-18
b3
y3
M-36
M
b4-18
b4
y4
y5
y6
b6

H

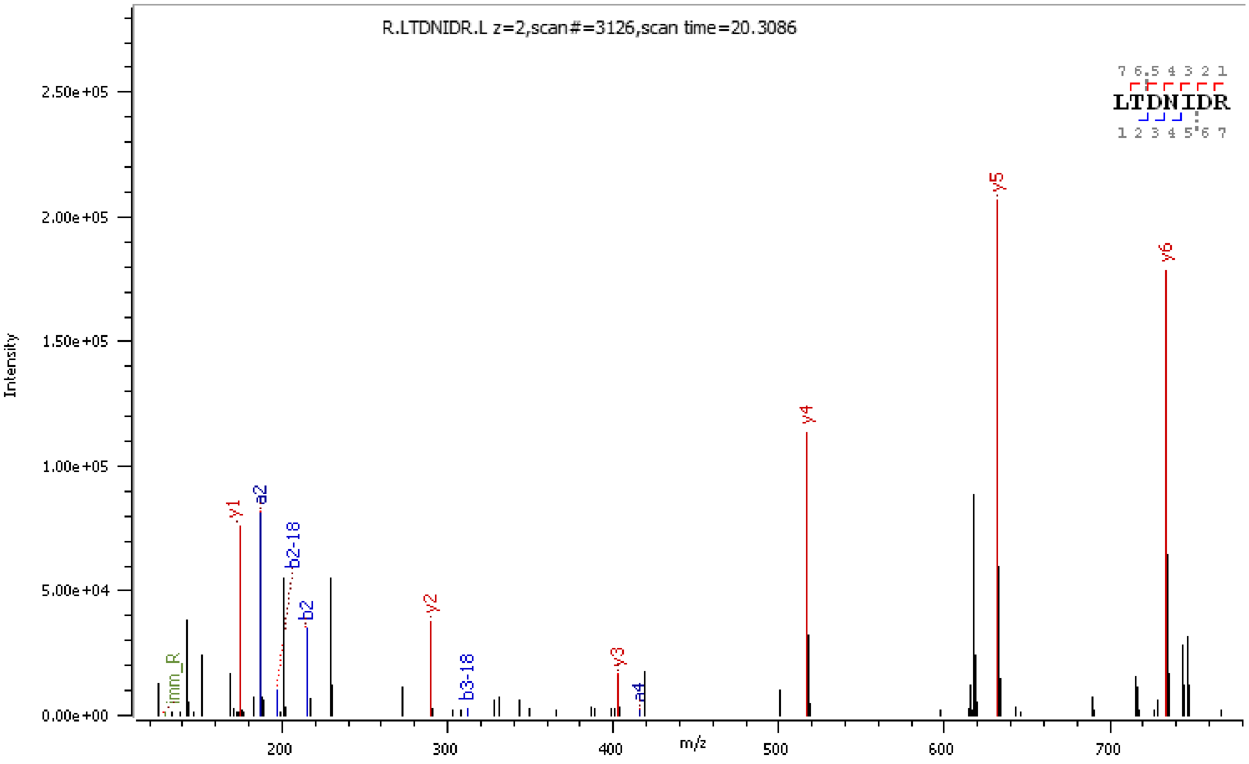
R.LTDNIDR.L z=2,scan#=3126,scan time=20.3086
LTDNIDR
Intensity
m/z
imm_R
y1
a2
b2-18
b2
y2
b3-18
y3
a4
y4
y5
y6

I

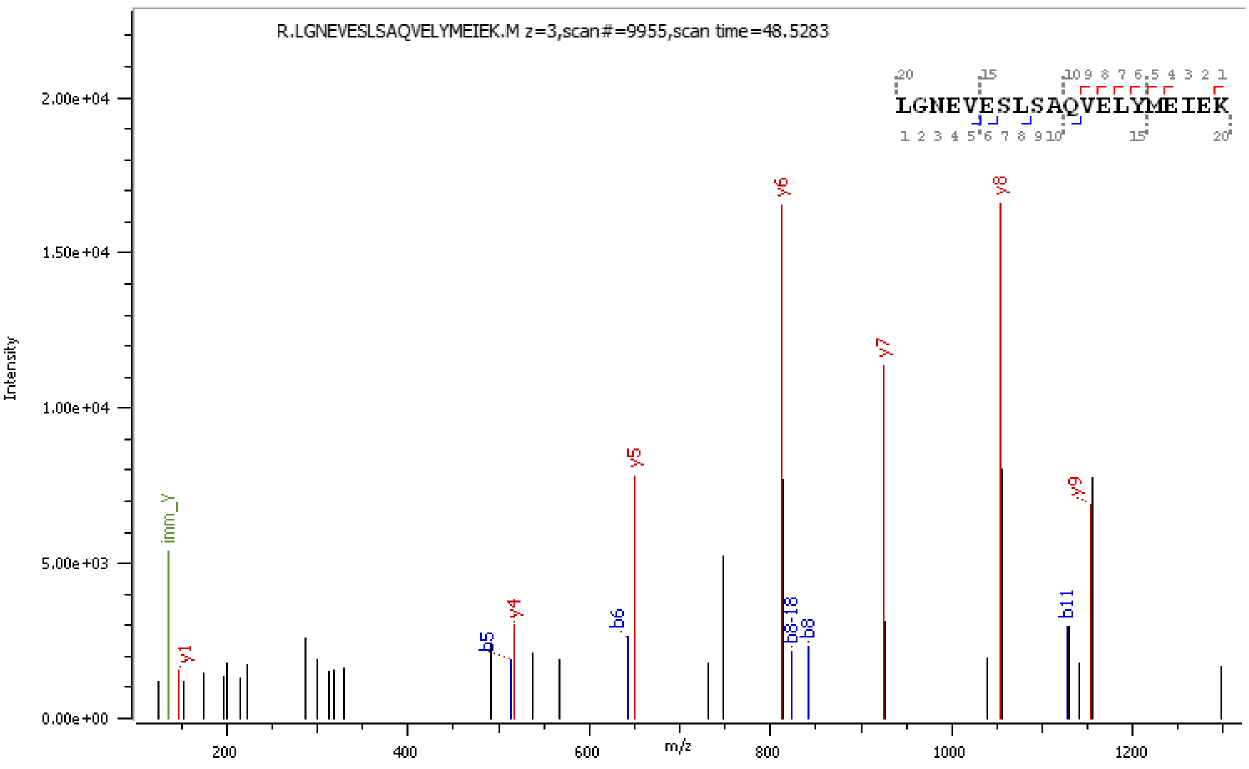
R.LGNEVESLSAQVELYMEIEK.M z=3,scan#=9955,scan time=48.5283
LGNEVESLSAQVELYMEIEK
Intensity
m/z
y1
imm_Y
b5
y4
b6
y5
y6
b8-18
b8
y7
y8
b11
y9

J

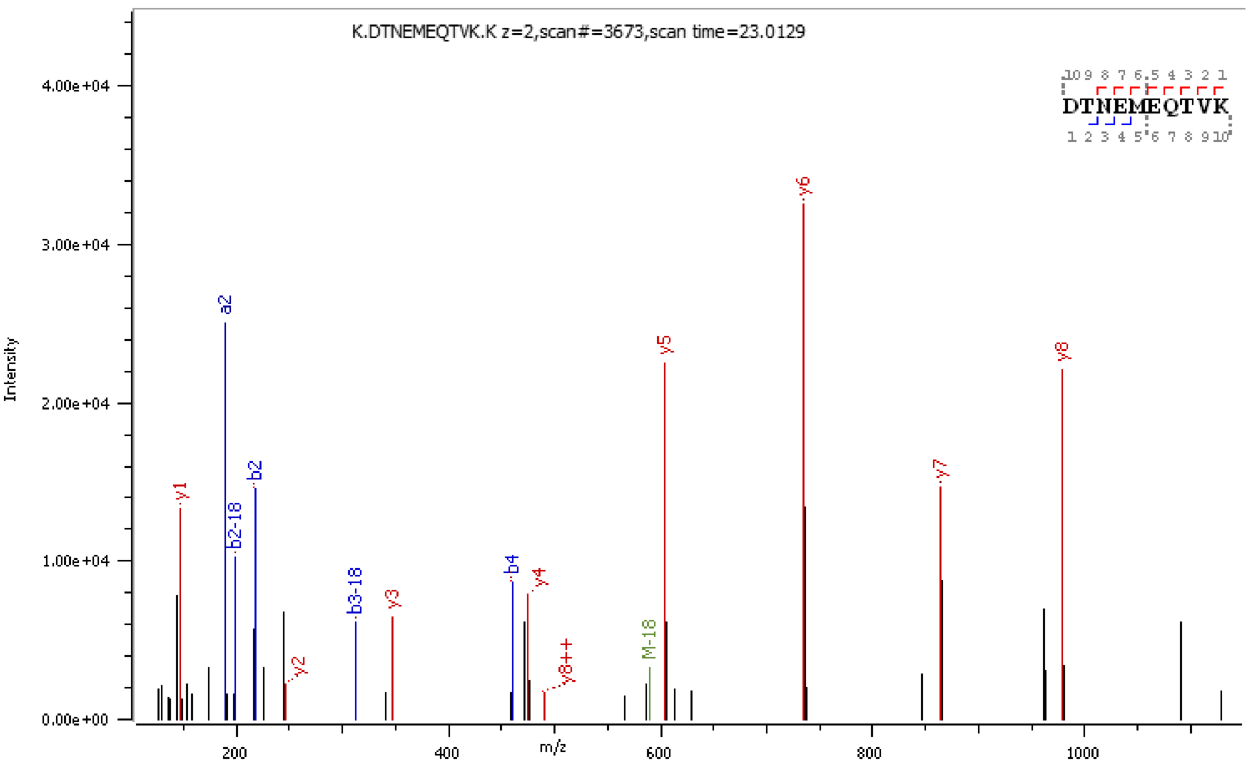
K.DTNEMEQTVK.K z=2,scan#=3673,scan time=23.0129
DTNEMEQTVK
Intensity
m/z
y1
a2
b2-18
b2
y2
b3-18
y3
b4
y4
y8++
M-18
y5
y6
y7
y8

**K**

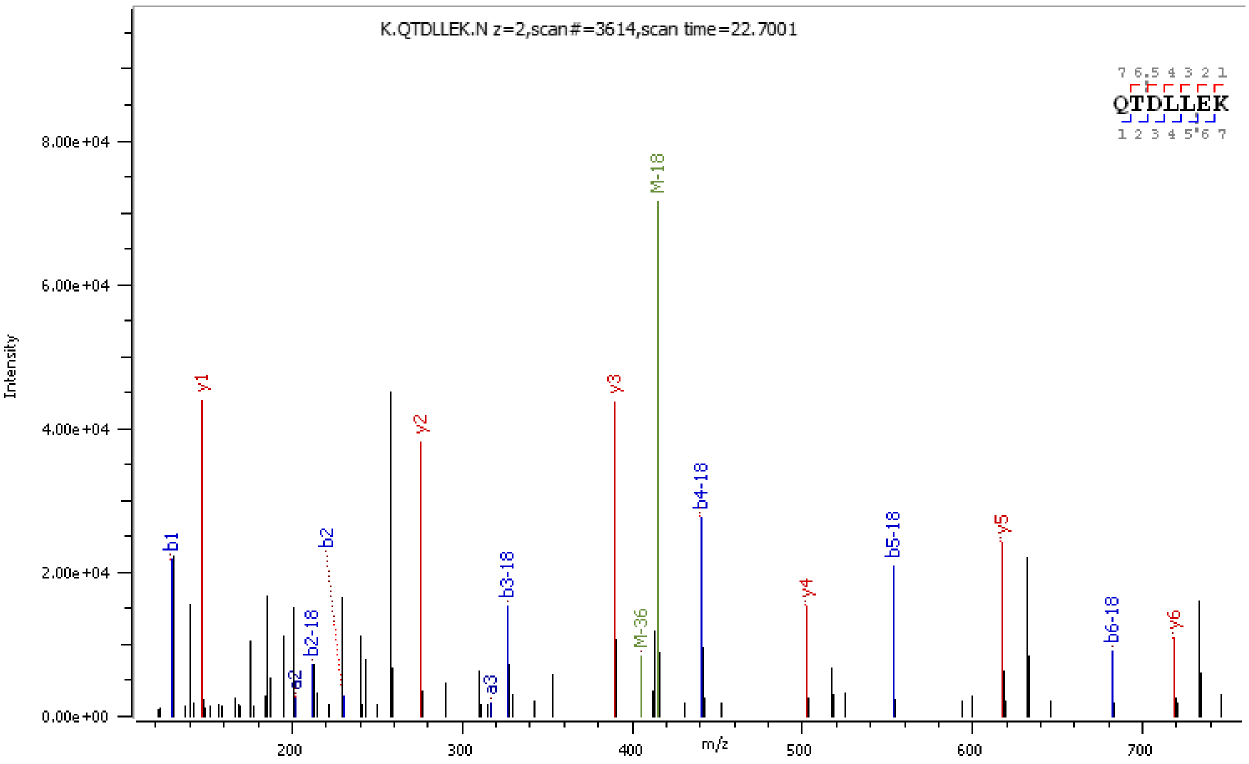
K.QTDLLEK.N z=2,scan#=3614,scan time=22.7001
QTDLLEK
Intensity
m/z
b1
y1
b2
a2
b2-18
y2
a3
b3-18
y3
M-36
M-18
b4-18
y4
b5-18
y5
b6-18
y6

**L**

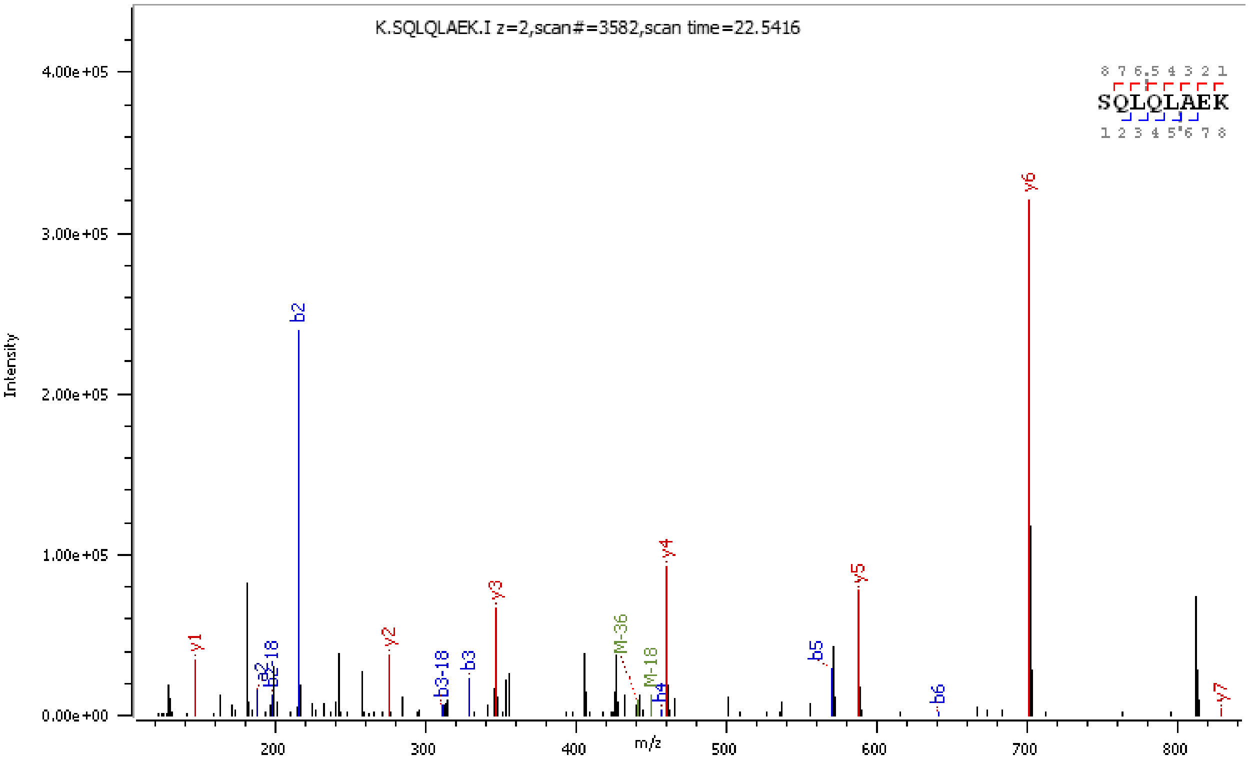
K.SQLQLAEK.I z=2,scan#=3582,scan time=22.5416
SQLQLAEK
Intensity
m/z
y1
b2
y2
b3-18
b3
y3
M-36
M-18
y4
b5
y5
b6
y6
y7

M

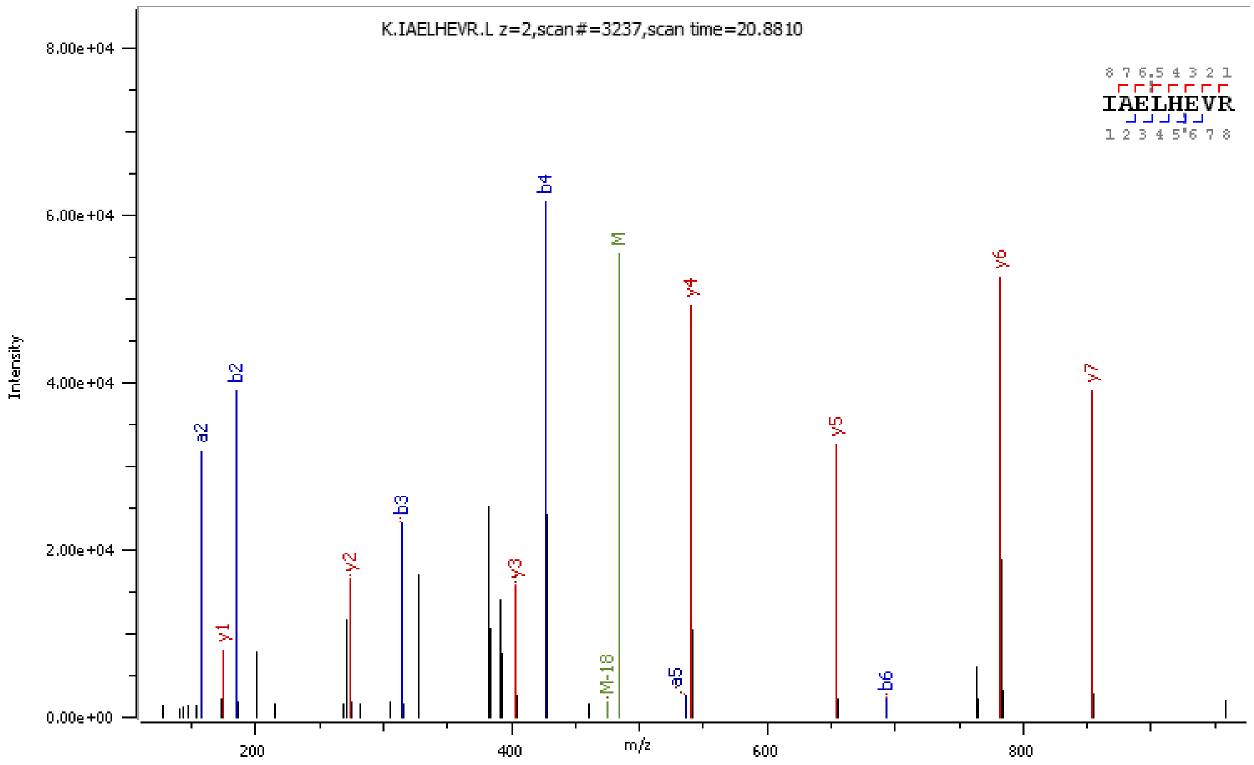
K.IAELHEVR.L z=2,scan#=3237,scan time=20.8810
IAELHEVR
Intensity
m/z
8.00e+04
6.00e+04
4.00e+04
2.00e+04
0.00e+00
200
400
600
800
a2
b2
y1
y2
b3
y3
b4
M-18
M
a5
y4
y5
b6
y6
y7

N

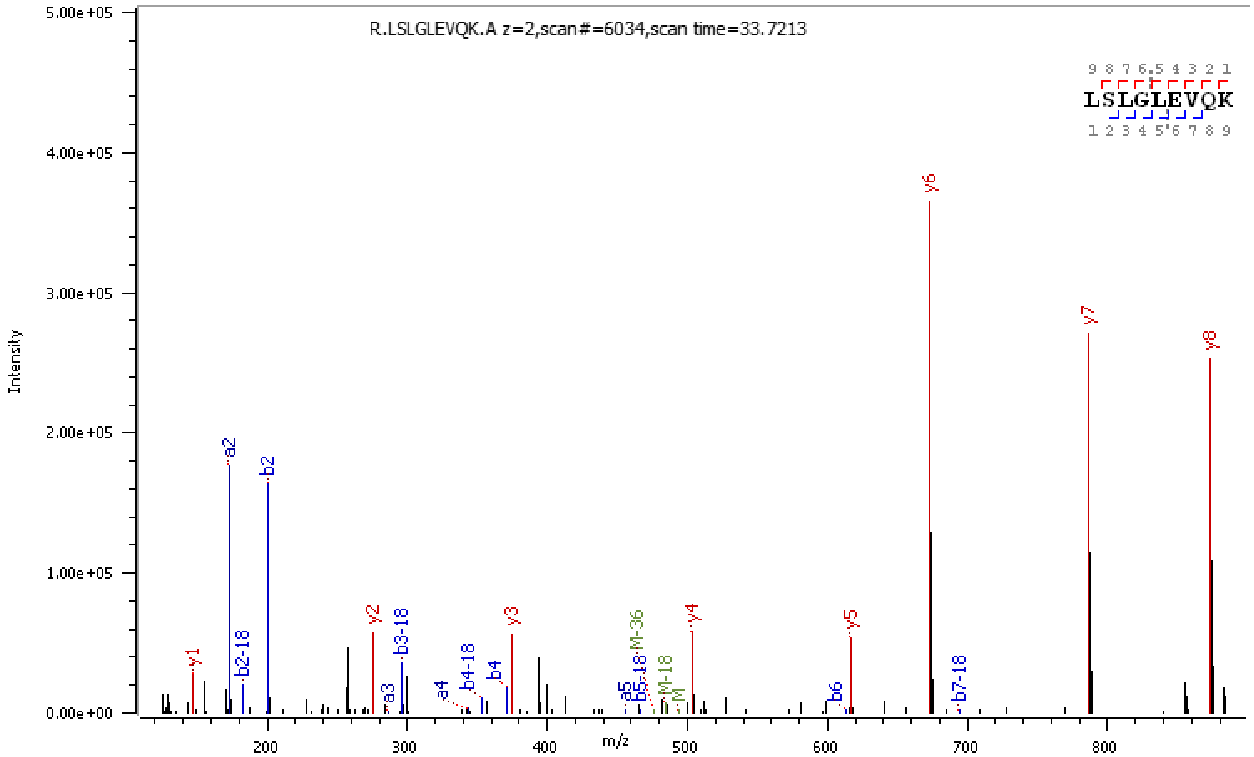
R.LSLGLEVQK.A z=2,scan#=6034,scan time=33.7213
LSLGLEVQK
Intensity
m/z
5.00e+05
4.00e+05
3.00e+05
2.00e+05
1.00e+05
0.00e+00
200
300
400
500
600
700
800
y1
a2
b2-18
b2
y2
a3
b3-18
a4
b4-18
b4
y3
a5
b5-18
M-36
M-18
M
y4
b6
y5
y6
b7-18
y7
y8

**O**

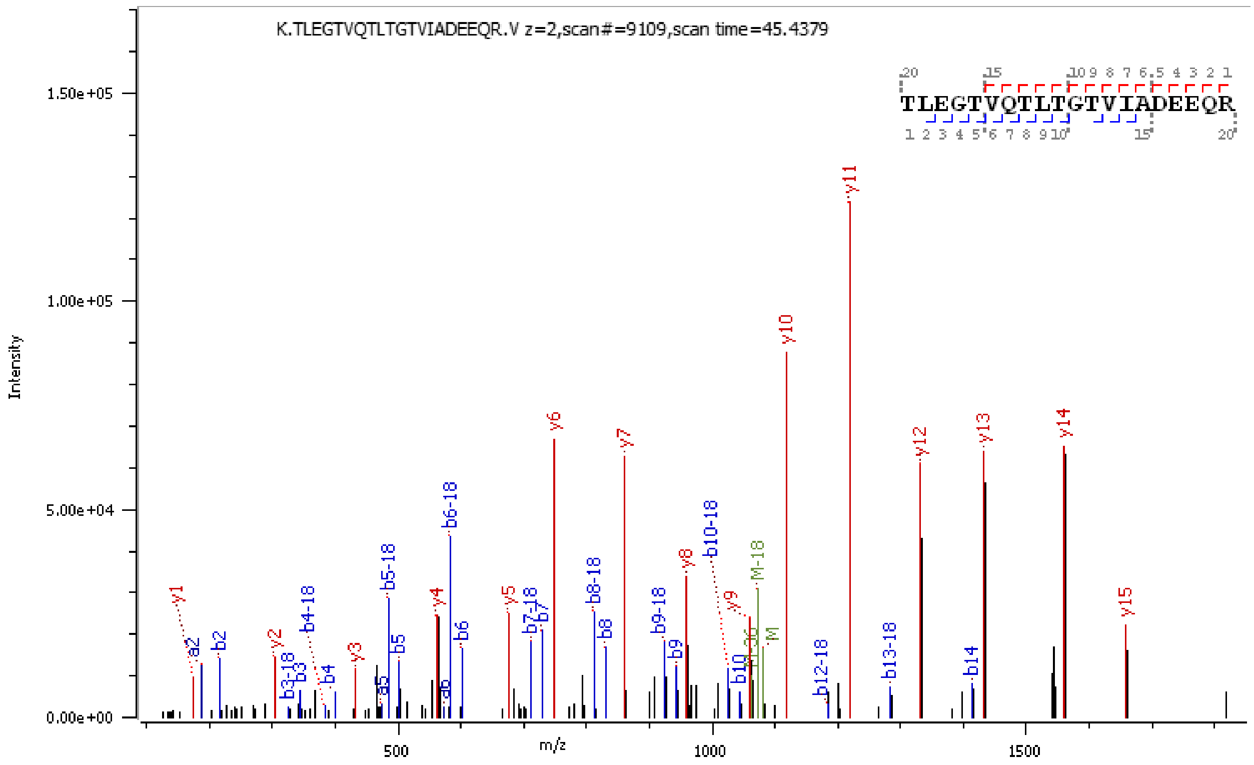

K.TLEGTVQTLTGTVIADEEQR.V z=2,scan#=9109,scan time=45.4379
TLEGTVQTLTGTVIADEEQR
Intensity
m/z


**P**

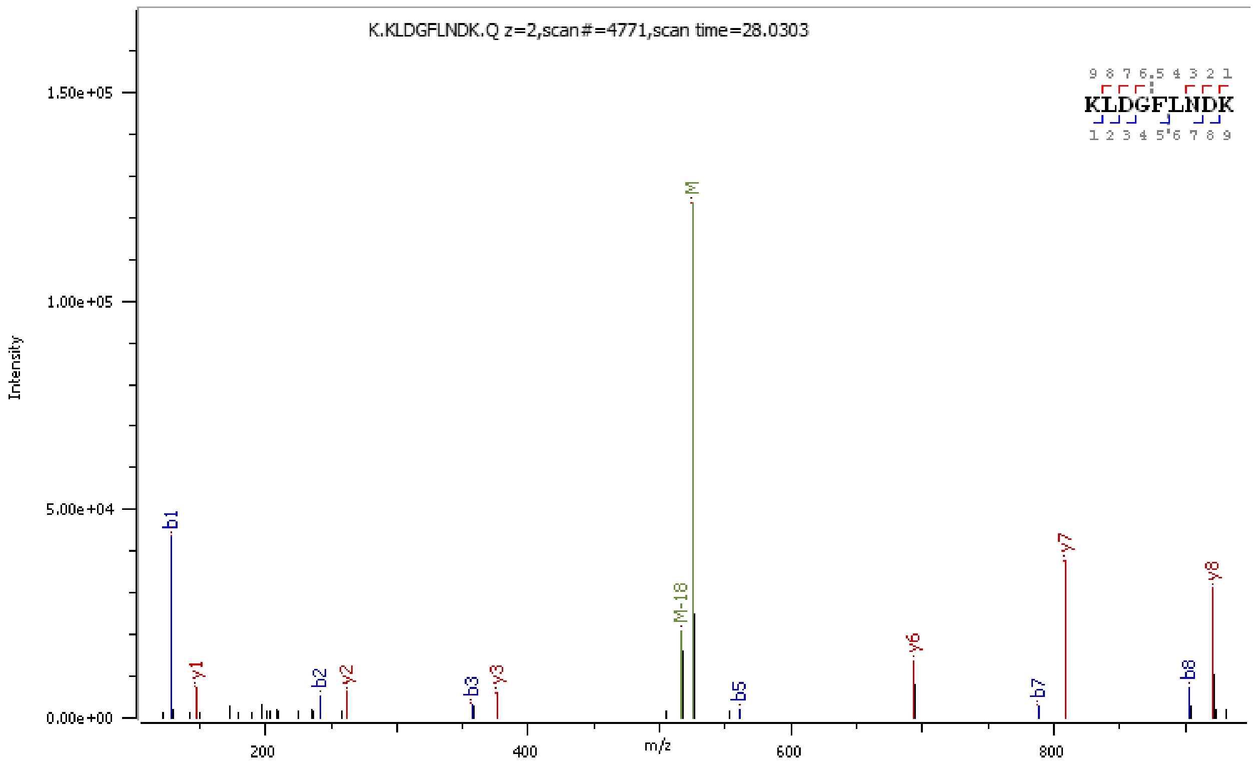

K.KLDGFLNDK.Q z=2,scan#=4771,scan time=28.0303
KLDGFLNDK
Intensity
m/z

**Q**

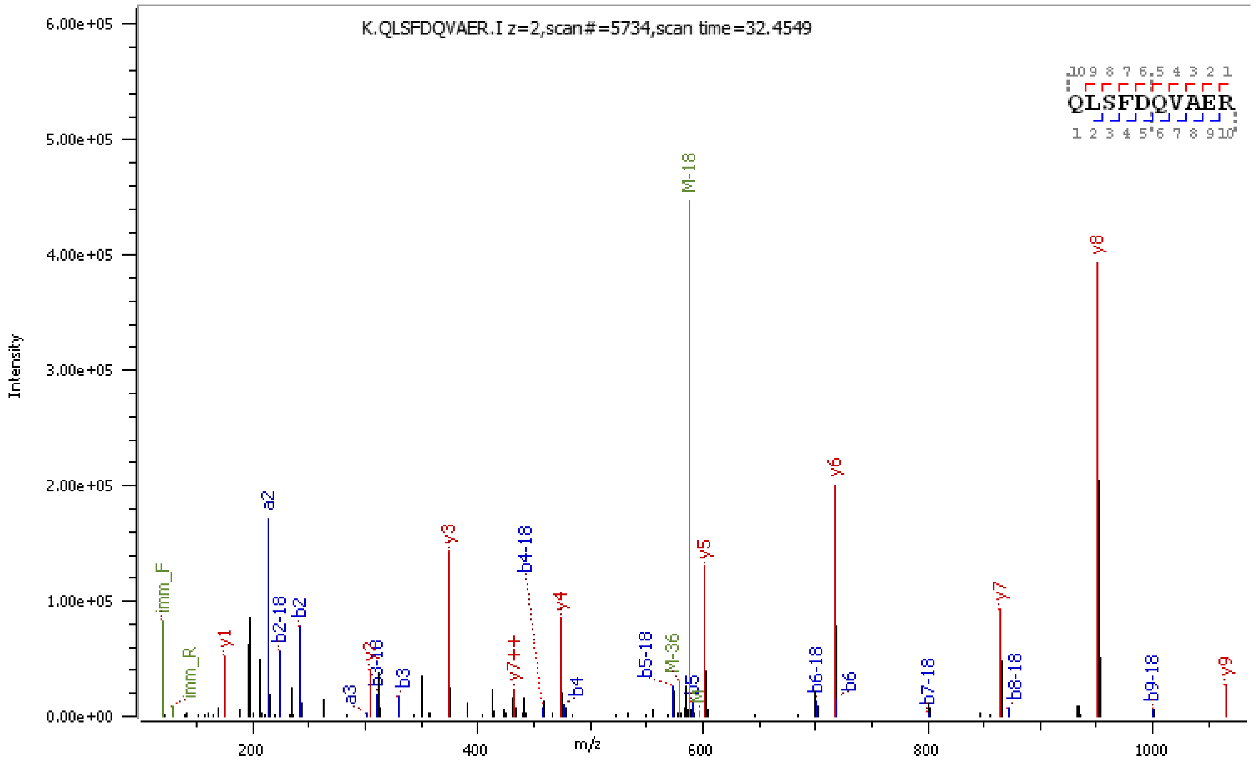
K.QLSFDQVAER.I z=2,scan#=5734,scan time=32.4549
QLSFDQVAER
Intensity
m/z
M-18
y8
y6
a2
y3
b4-18
y5
y4
y7
b2-18
b2
y1
imm_F
imm_R
y7++
b5-18
M-36
b6-18
b6
b7-18
b8-18
b9-18
y9
a3
b3
b4
b5

**R**

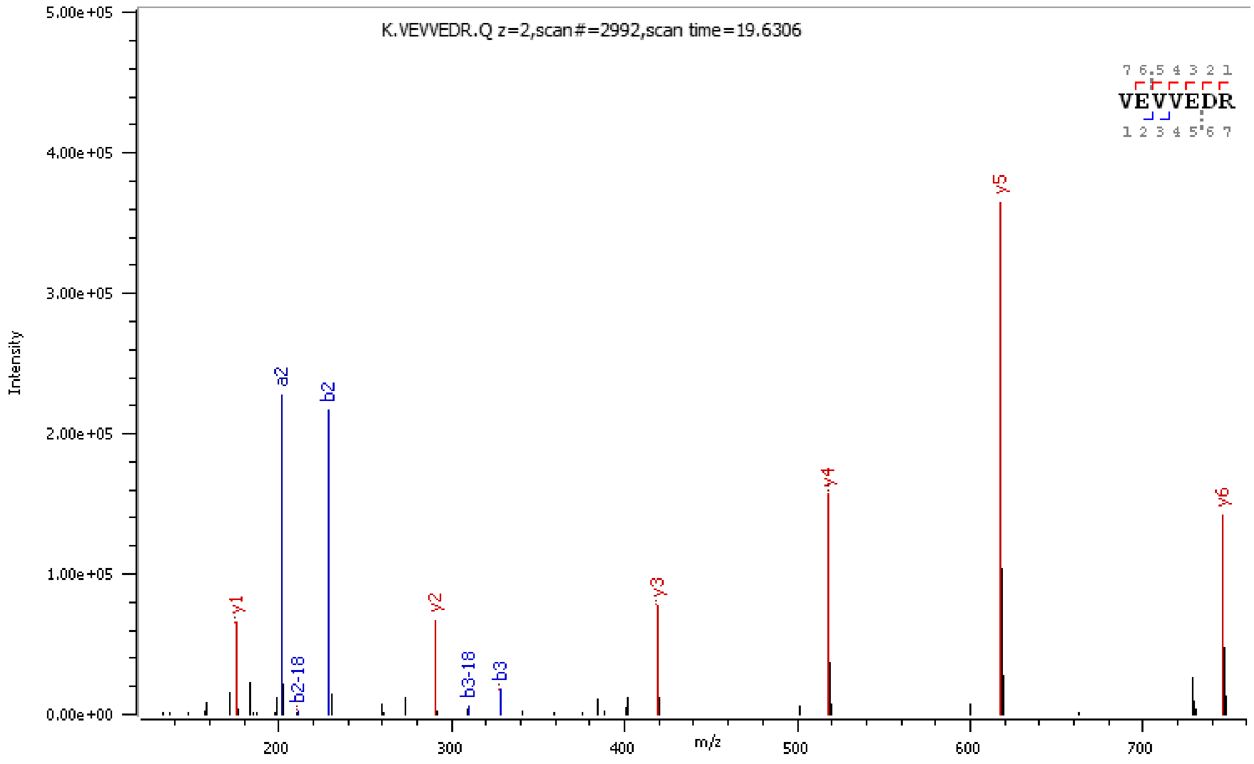
K.VEVVEDR.Q z=2,scan#=2992,scan time=19.6306
VEVVEDR
Intensity
m/z
y5
a2
b2
y4
y6
y3
y1
y2
b2-18
b3-18
b3

### Figure S3

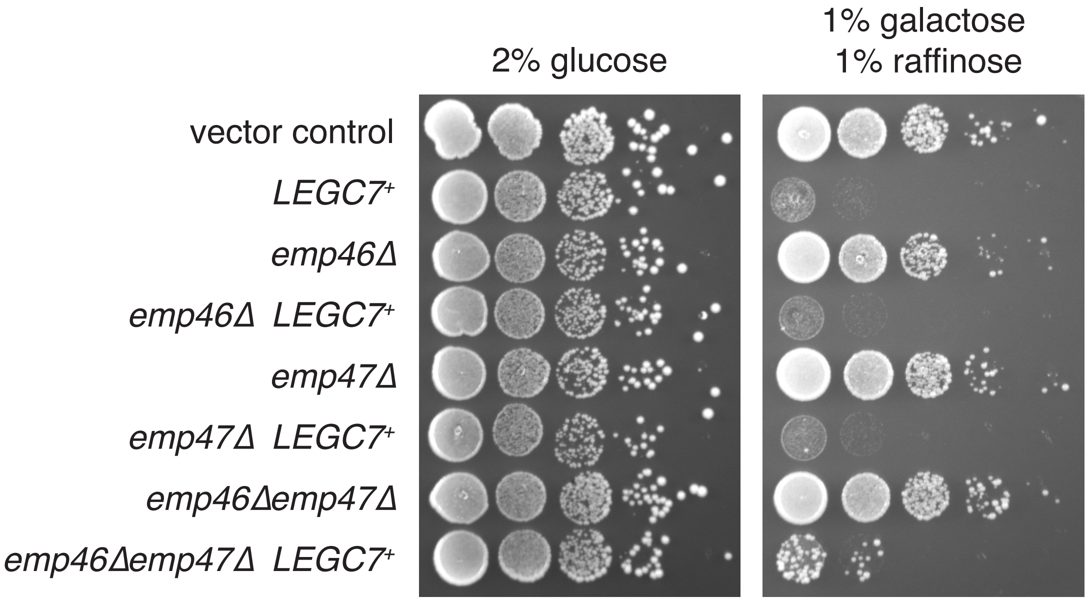

2% glucose
1% galactose
1% raffinose
vector control
LEGC7+
emp46Δ
emp46Δ LEGC7+
emp47Δ
emp47Δ LEGC7+
emp46Δemp47Δ
emp46Δemp47Δ LEGC7+

### Figure S4

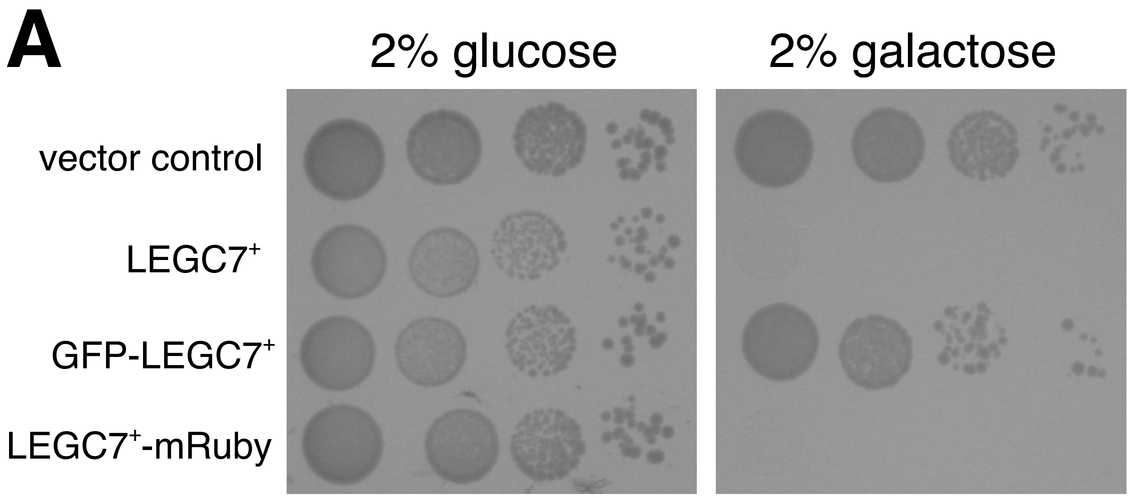
A
2% glucose
2% galactose
vector control
LEGC7$^{+}$
GFP-LEGC7$^{+}$
LEGC7$^{+}$-mRuby

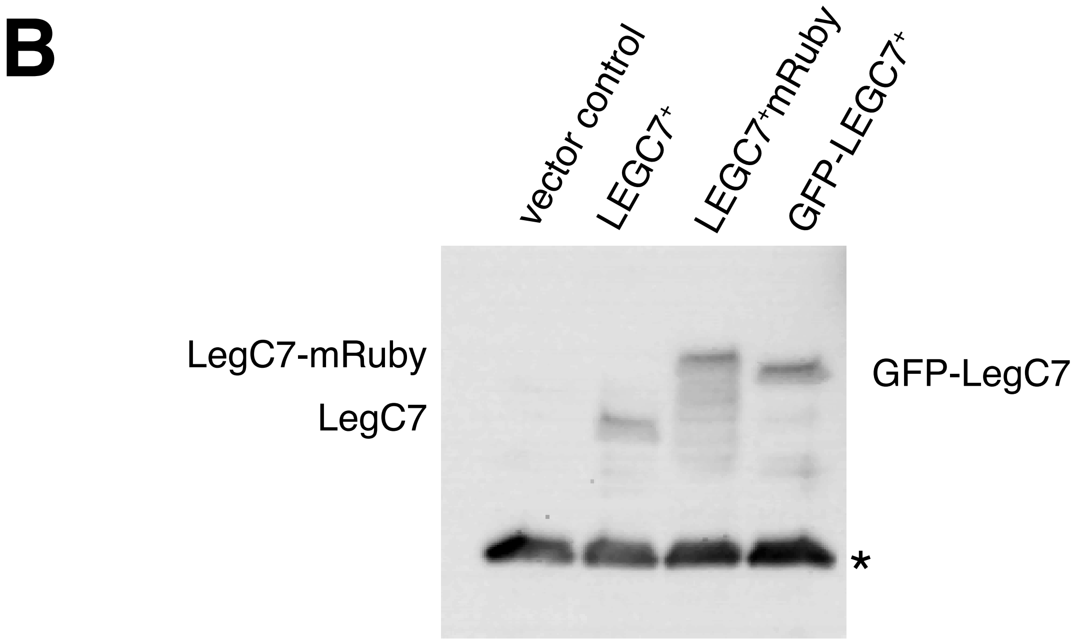
B
vector control
LEGC7$^{+}$
LEGC7$^{+}$mRuby
GFP-LEGC7$^{+}$
LegC7-mRuby
LegC7
GFP-LegC7
*

### Figure S5

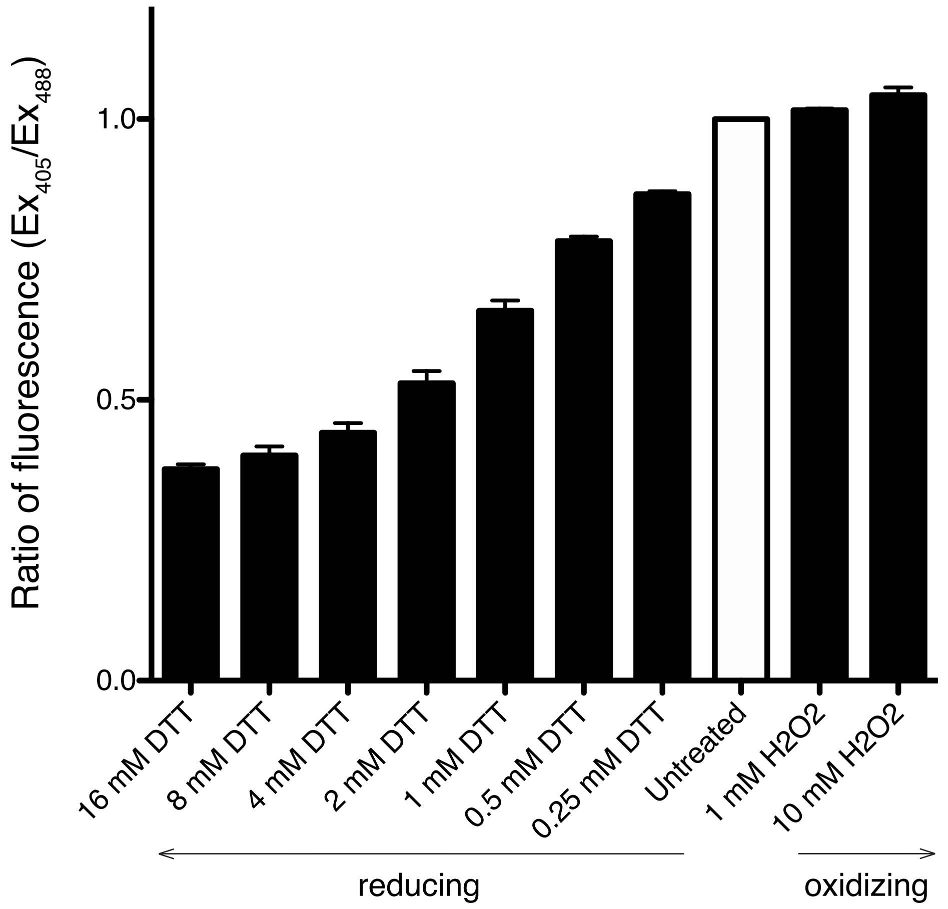

Ratio of fluorescence ($Ex_{405}/Ex_{488}$)
1.0
0.5
0.0
16 mM DTT
8 mM DTT
4 mM DTT
2 mM DTT
1 mM DTT
0.5 mM DTT
0.25 mM DTT
Untreated
1 mM H2O2
10 mM H2O2
reducing
oxidizing

### Figure S6

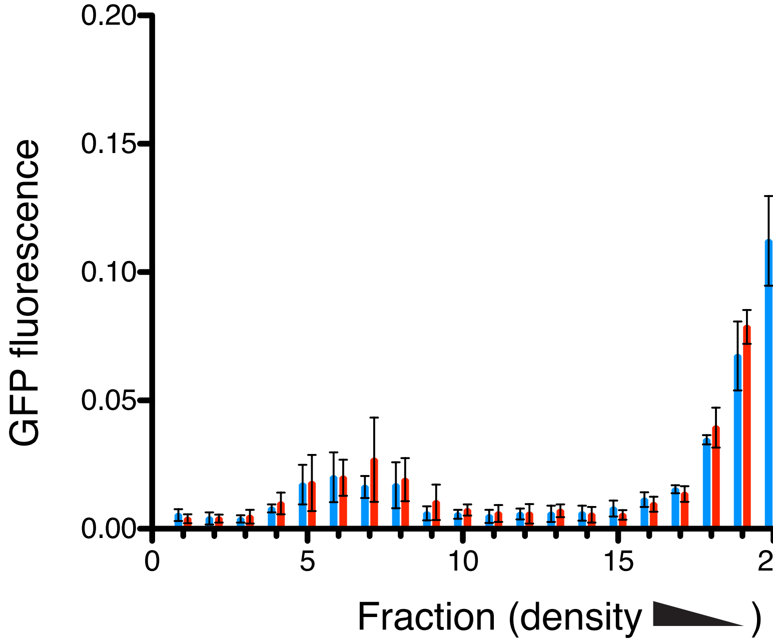
GFP fluorescence
0.20
0.15
0.10
0.05
0.00
0
5
10
15
20
25
Fraction (density )
